# Population dynamics in extreme environments: shared histories of strong selection lead to predictable patterns across separate high-elevation populations of small mammals

**DOI:** 10.64898/2026.11.06.764945

**Authors:** Nathan R. Senner, Brett R. Addis, Ryan Mahar, Thomas C. Nelson, K. Trey Sasser, Rena M. Schweizer, Sarah R. Senese, Maria Stager, Jonathan P. Velotta, Luke R. Wilde, Cole J. Wolf, Zachary A. Cheviron

## Abstract

Many frameworks exist to identify species that are vulnerable to environmental change, but we still generally lack the ability to anticipate species’ responses when changes occur. Exploring patterns of trait evolution along environmental gradients could help this process: conditions become more predictable and less variable at the ends of gradients, with past studies hypothesizing that consistently harsh environments will result in predictable evolutionary patterns. To test this hypothesis, we characterized genetic and phenotypic variation, as well as selection on this variation, in deer mice (*Peromyscus maniculatus*) along two transects spanning from 2120-4200m in the Colorado Rocky Mountains, U.S.A. We found that both genetic and phenotypic variation declined with elevation, that high elevation populations had higher rates of survival, and that fewer traits influenced survival at high elevations. These results suggest extreme environments may lead to predictable consequences for populations and provide a basis for forecasting population vulnerability to environmental change.

## INTRODUCTION

One of the core goals of evolutionary ecology is to determine the degree to which population dynamics and evolutionary change follow predictable patterns (Mas et al., 2020; Reiskind et al., 2021). The development of accurate predictive eco-evolutionary frameworks would aid both our understanding of evolutionary processes and our ability to identify which populations are most vulnerable to future environmental changes. As a result, myriad attempts have been made to link a population’s evolutionary history (Wiens & Graham, 2005), trait architecture (Stern & Orgogozo, 2008), physiological pathways (Taborsky et al., 2021), biogeography (Losos & Ricklefs, 2009), body size (Lanfear et al., 2014), and biotic interactions (Ickin et al., 2024) with their future evolutionary trajectory. These efforts, however, have met with varying degrees of success (Bitter et al., 2021). As a result, we still lack the predictive power to anticipate the effects of environmental change on most species, despite the importance of such predictions in potentially slowing the loss of biodiversity (Cahill et al., 2013).

Broadly characterizing patterns of environmental variation and its predictability may be key to the development of accurate eco-evolutionary predictive frameworks. These two factors are widely recognized as playing important roles in shaping evolutionary processes and are thought to strongly affect the types of traits likely to evolve (Botero et al., 2015; Gabriel et al., 2005; Stager et al., 2025), as well as the types of interactions exhibited between species (Bertness & Callaway, 1994), in a given environment. The more predictable and less variable the environment, the more likely fitness is to be determined by survival rather than reproduction, and the more likely there is to be a stable fitness optimum associated with a small number of survival related traits. In these environments, directional selection should then push populations toward that single fitness optimum, followed by stabilizing selection that should act to keep the population at the optimum once it has been reached (Saether & Bakke, 2000; Smith & Fretwell, 1974). Over time, low variability and high predictability environments are thus predicted to winnow a population’s genetic and phenotypic variation *via* purifying selection at fitness-related loci and the regions linked with them across the genome (Leiby & Marx, 2014; Zhang, 2012). Once locally adapted to invariable and highly predictable environments, populations may lack the genetic or phenotypic variation to respond to novel environmental conditions, leading to rapid population declines when environmental changes do occur (Studds et al., 2017). Quantifying gradients in environmental predictability and variability could therefore provide an avenue to identify the degree to which eco-evolutionary dynamics can be broadly predictable across species and geographic scales.

The extreme environments that exist at the ends of abiotic spectra – such as temperature, salinity, or oxygen availability – provide the ideal opportunity to test these predictions (Conklin et al., 2017). Extreme environments are characterized by consistently harsh abiotic conditions and tend to support relatively simple biotic communities (Stevens & Tello, 2018). The species that do occupy extreme environments often exhibit novel traits (e.g., hemoglobin loss in Antarctic fishes; Malekar *et al*. 2018). These novel traits, in turn, likely reflect both a history of strong directional selection and the existence of absolute physiological limits past which further adaptation is impossible, constraining the evolutionary potential of extremophile populations (Somero, 2010). Accordingly, studies of the genetic architecture of the novel traits of extremophiles indicate that only a few loci may encode them and that the traits themselves may be subject to antagonistic pleiotropy linked with strong trade-offs that further limit the ability of extremophiles to adapt to environmental change (Gerstein et al., 2012, 2015; Szűcs et al., 2017). Extreme environments are thus expected to lead to especially predictable patterns of evolutionary change (Greenway et al., 2020; Tobler et al., 2011), meaning that populations living in environments along a spectrum of extremeness may exhibit a continuum of evolutionary predictability.

Among the most well characterized extreme environments are those found at high elevations, where chronic hypoxia has led to a suite of novel adaptations in a number of vertebrate lineages (Storz et al., 2010, 2019; Storz & Scott, 2019). In comparison with low-elevation populations, high-elevation populations of deer mice (*Peromyscus maniculatus*), for instance, exhibit a specialized high-elevation hemoglobin variant that has higher oxygen binding affinity (Storz et al., 2009), pulmonary adaptations to ensure a higher arterial O_2_ saturation under hypoxia (Tate et al., 2017, 2020), a shift to more efficient fuel usage while exercising (Lau et al., 2017), greater lipid oxidation rate during thermogenesis (Cheviron et al., 2012; Cheviron, Natarajan, et al., 2014), higher oxidative fiber density and increased capillarity in muscles (Scott et al., 2015), and larger left heart ventricles, which supply blood flow for systemic circulation (Velotta et al., 2018). Past work suggests that some of these traits have experienced both historical and contemporary selection (Wilde et al., 2019), and that these traits correspond with a suite of genetic changes (Schweizer et al., 2019; Storz et al., 2009, 2012). Nonetheless, there is little evidence indicating whether genetic variation, phenotypic variation, and population dynamics vary predictably across the deer mouse elevational range (Schweizer et al., 2021), which extends to environments below sea level (Hock, 1964).

To assess the degree to which environmental gradients may produce predictable patterns in the demographic and evolutionary histories of the populations distributed along them, we set up replicated elevational transects in the Rocky Mountains of Colorado, USA, along which we characterized spatial patterns of phenotypic and genetic variation in deer mice, as well as the strength of contemporary selection pressures on traits known to influence survival across the deer mouse range. We hypothesized that populations occupying similar elevations along each transect would exhibit similar amounts of phenotypic and genetic variation, as well as experience similar strengths of selection on similar traits. We then predicted higher elevation populations would experience stronger selection, resulting in lower genetic and phenotypic variation, and fewer traits under contemporary selection, with those traits under selection being related to the ability of deer mice to withstand the effects of hypoxia. Taken together, our study will aid efforts to develop predictive eco-evolutionary frameworks that can forecast how species at specific points along environmental gradients may be able to respond to environmental change.

## METHODS

### Study System

Deer mice have the largest elevational range of any North American mammal and can be found from deserts below sea level to the tops of the tallest Rocky Mountains (Hock, 1964). Across this range, deer mice have repeatedly exhibited local adaptations to a range of habitats, including low-elevation sandhill grasslands (Barrett et al., 2018; Pfeifer et al., 2017) and high-elevation tundra (Chappell et al., 2007; Hayes & O’Connor, 1999). The combination of a large elevational range and repeated local adaptation to different environments implies that deer mice have a considerable ability to both disperse geographically and respond to novel environments, making them an excellent study species with which to assess the predictability of population dynamics under different environmental conditions.

To monitor deer mice across a portion of their elevational range, we established two elevational transects in the Front Range of the Colorado Rocky Mountains in the western U.S.A., each spanning from 2,120-4,200 meters above sea level – a southern transect along the Clear Creek watershed through Arapahoe National Forest (2016-2018) and a northern transect along the Big Thompson watershed through Roosevelt National Forest and Rocky Mountain National Park (2017-2018; Fig. 1). Along each transect, we established three plots at which we sampled mice twice each year – once immediately after snowmelt in spring (Apr-Jun) and once prior to snow coverage in fall (Sep-Oct). Given that deer mice are habitat generalists (Williams, 1955), each plot differed in its habitat type, but all were placed within open habitats predominantly characterized by herbaceous vegetation.

**Figure 1.**
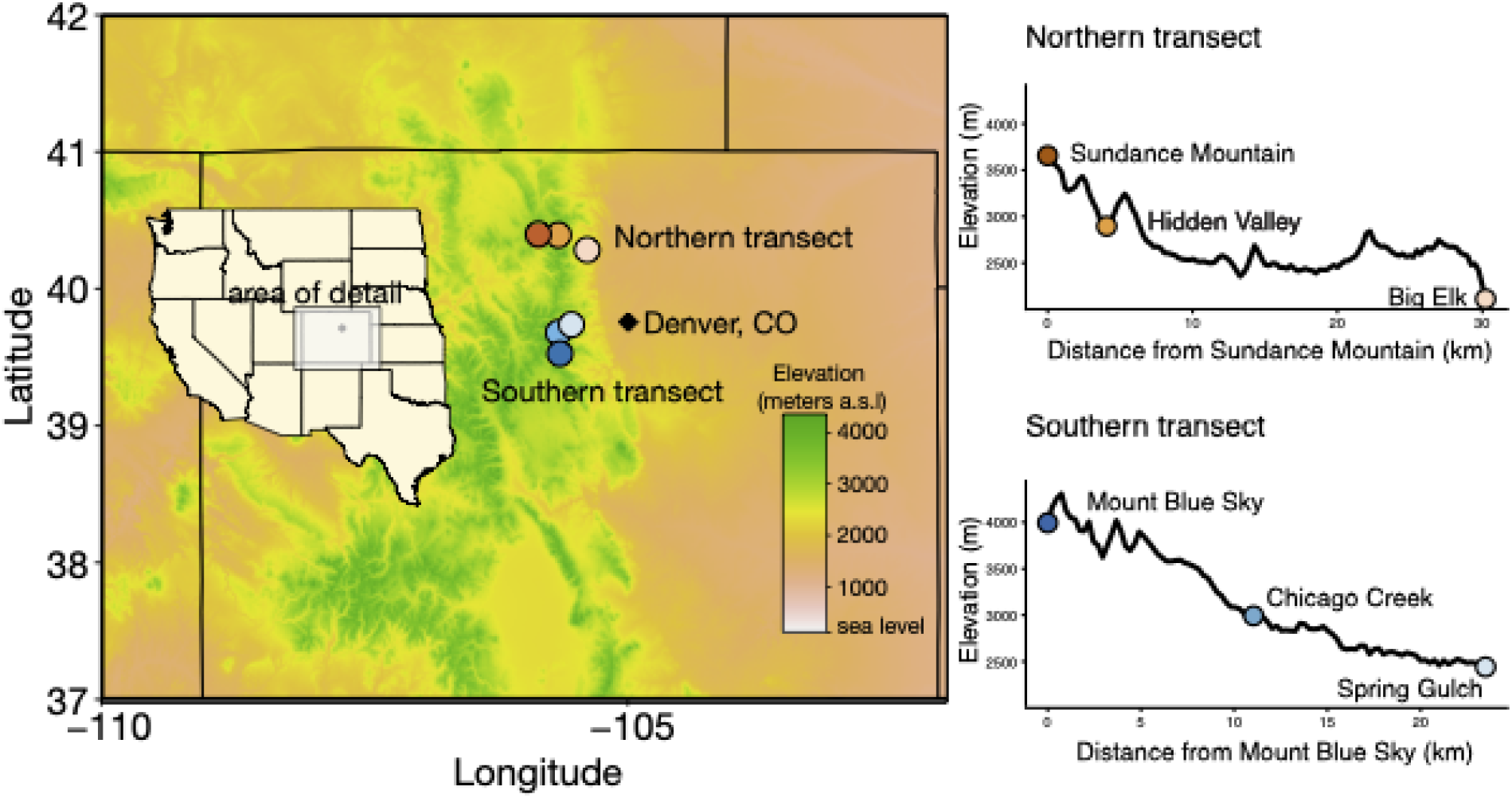
Map of study area. On left, geographic representation of demographic sites, with each site represented by a colored circle overlaid on a topographic map. On right, the elevation and ground-based distances of the demographic sites along the northern and southern transects.

### Field Sampling

Each demographic plot was comprised of a rectangular grid of Sherman live traps spaced 10m apart and ranging in size from 2-4ha, depending on the amount of available habitat. During each sampling period, we trapped at a site for 3-7 nights with appropriate weather or until at least 30 unique individuals had been captured and phenotyped (see below). Each trap was set between 15:00-19:00, baited with peanut butter and oats, and filled with polyester fiber. We checked traps each morning between 07:00-09:00 and immediately released non target species; all traps then remained closed until >15:00.

### Phenotyping

We chose to measure a suite of morphological and physiological phenotypes that have previously been linked with deer mouse or small rodent fitness: (1) anogenital distance, a proxy for testosterone levels in males (Godsall et al., 2014); (2) head width and (3) length, which are associated with bite force and diet (Williams et al., 2009); (4) tarsal-metatarsal length, which contributes to running speed (Andrew et al., 2016; Djawdan & Garland, 1988); (5) body condition, measured as an individual’s residual from a regression of body mass on tarsus length, which can influence their ability to withstand food shortages (Wolf, 2023); (6) blood hemoglobin concentration and (7) hematocrit level, which correlate with blood-oxygen carrying capacity and aerobic performance (Lui et al., 2015; Tate et al., 2020); (8) resting metabolic rate (RMR), a proxy for an individual’s baseline energetic requirements (Speakman et al., 2003); (9) cold-induced summit metabolic rate (M_sum_), which represents an individual’s maximum aerobic capacity; and, (10) metabolic scope (i.e., the difference between M_sum_ and RMR), which represents the ability of an individual to up-regulate its aerobic output to meet a thermogenic challenge (Cheviron, Connaty, et al., 2014; Hayes & O’Connor, 1999; Wilde et al., 2019).

For each new mouse captured, we recorded the individual’s anogenital distance (females: ≤4 mm), mass, and age class (juveniles: ≤16 g). We inserted a 12.5-mm, 134.2-kHz PIT tag (HPT 12, Biomark, Boise, ID, USA) into their interscapular region for identification during subsequent trapping bouts. We also collected an ear clip as a genetic sample and a ≤50-μl blood sample from the maxillary vein for measures of hemoglobin and hematocrit. We quantified an individual’s hemoglobin concentration with a HemoCue Hb 201+ analyzer (HemoCue AB, Angelholm, Sweden) and their hematocrit value by measuring the column height of packed red blood cells relative to the whole blood column height in a heparinized capillary tube following centrifugation at 12,000rpm for 5 minutes. Finally, we measured each morphological trait to the nearest millimeter using hand-held calipers.

We measured an individual’s RMR at 27°C and M_sum_ at -5°C within a few hours of capture (either the initial or a subsequent capture) using open-flow respirometry following established protocols optimized for deer mice (Cheviron *et al*. 2014; Wilde *et al*. 2019; see Supplementary Material for more details). We recorded the oxygen consumption of each individual in the RMR trials over a series of 15 min intervals allowing for alternation of measurements between multiple animals and an empty, baseline chamber. This process was repeated until a stable measure – a 10 min period with no visually significant change in slope of oxygen consumption – was collected for each individual (1-4 × 15 min intervals). We calculated M_sum_ as the maximum rate of oxygen consumption averaged over a continuous 3 min period during a 30-min trial. We calculated both RMR and M_sum_ for each individual using customized scripts (Stager et al., 2020) with equations from Lighton (2008) and then corrected those values for the individual’s body mass by dividing by their mass at the time of the metabolic trial.

We could not necessarily perform metabolic measurements on every mouse on the day they were first captured because of time constraints. For mice randomly chosen to not undergo a respirometry trial on the day of their first capture, their morphological measurements and body mass were taken soon after capture and then the mouse was released at its capture site; for individuals undergoing respirometry trials, the measurements were taken 30 minutes following the termination of their M_sum_ trial.

### Genetic Sample Processing

To assess the relatedness (see below) and potential for differences in levels of genetic variation among the mice from each transect and demographic plot, we extracted DNA from tissue samples collected from each mouse and then developed a custom library of single nucleotide polymorphisms (SNPs). We extracted the DNA using a Qiagen DNeasy Blood and Tissue Extraction Kit (Valencia, CA, USA) and then developed our SNP libraries with an NEBNext Ultra DNA Library Prep Kit (Ipswich, MA, USA) using the double digest restriction site associated capture protocol developed by Ali *et al*. (2016). Briefly, we digested whole genomic DNA with two restriction enzymes (*BfaI* and *PstI*), ligated the resulting fragments with uniquely barcoded adaptor sequences for each individual, performed a PCR amplification, and size selected ∼400 bp fragments using a Pippin Prep (Sage Science, Beverly, MA, USA). Libraries were sequenced at the University of California, Berkeley Sequencing Facility (Berkeley, CA, USA) on three lanes of an Illumina HiSeq 4000 platform, generating 603 million, 150-bp paired-end reads.

To identify SNPs, we demultiplexed our resulting reads and removed true PCR duplicates using custom Python scripts (Kolis et al., 2022). This resulted in 543,776,631 reads retained. We mapped our reads to the deer mouse genome (ftp://ftp.ncbi.nih.gov/genomes/Peromyscus_maniculatus_bairdii/GFF/) using BWA MEM (Li & Durbin, 2009) and indexed them with SAMtools (Li et al., 2009), retaining 89.3% of our original reads. We then called SNPs using the STACKS pipeline (v2.62; Catchen et al., 2011, 2013) for aligned sequence data. Following this, we applied a series of additional filtering steps in VCFtools (v0.1.17; Danececk et al. 2011): First, we removed three duplicate individuals and any monomorphic sites, leaving 2,183,359 candidate SNPs across 486 individuals. We then applied an iterative filtering procedure that alternated individual- and site-level missingness thresholds. Specifically, we **(1)** removed individuals with >80% missing genotypes (486 to 339 individuals); **(2)** filtered sites to a minor allele frequency of ≥0.10 and minimum mean depth ≥5x (2,183,359 to 9,225 sites); **(3)** required site-level genotyping in ≥50% of individuals (9,225 to 9,044 sites); **(4)** removed individuals with >30% missing genotypes (339 to 196 individuals); **(5)** required site-level genotyping in ≥30% of individuals (i.e., max-missingness of 70%; 9,044 to 8,020 sites); **(6)** removed individuals with >25% missing genotypes (196 to 180 individuals); and **(7)** applied a final site-completeness filter requiring genotyping in ≥5% of individuals (i.e., max-missingness of 95%), yielding our final dataset of 818 SNPs across 180 individuals (see full step-by-step counts and *posthoc* quality checks in the Supplementary Methods and Supplementary Table 1).

### Statistical Analyses

Our statistical analyses involved three separate steps to identify patterns along our elevational transects in **(1)** genetic variation, **(2)** phenotypic variation, and **(3)** survival and natural selection:

To assess the amount of genetic variation within each demographic plot and across elevations, as well as whether different populations exhibit different site frequency spectra – a potential signature of historical population bottlenecks – we used the program ‘VCFtools.’ This allowed us to estimate pairwise measures of F_st_ – a measure of population divergence – among sites, levels of heterozygosity for each individual at each of our 818 SNPs, and a separate site frequency spectrum for each demographic site derived from the minor allele frequency at each of our SNPs generated using the function *freq*. To determine whether levels of individual heterozygosity varied across elevations, we used a mixed effect beta regression in the ‘gamlss’ package (Stasinopoulos, 2024) in the R Programming Environment (v. 4.1.1; The R Foundation for Statistical Computing 2021), with observed heterozygosity as the dependent variable, elevation as the predictor variable, and transect as a random effect. To quantify whether our demographic sites had different site frequency spectra, we used Kolmogorov-Smirnov tests in the R package ‘dgof’ (Arnold & Emerson, 2011) to compare between each pair of sites, followed by a beta regression to determine whether the difference in elevation between two sites predicted the degree to which their site frequency spectra differed. We used a similar approach to assess whether elevation influenced variation in F_st_ except using a beta regression instead of a Kolmogorov-Smirnov test. In this case, the F_st_ between each pair of sites was the dependent variable, and the elevational difference between those sites was the predictor variable.

To characterize the amount of phenotypic variation in our 10 focal traits within each demographic plot and across elevations, we constrained our analyses to only phenotypic measures gathered from adult mice, as we could not precisely age juveniles. For three traits, we also limited our analyses to only males – anogenital distance, which is likely only under selection in males (Godsall *et al*. 2014), and mass and body condition, for which our measures in females were influenced by our inability to characterize active pregnancies in the field. We then estimated coefficients of variation – a unitless measure of variation robust to differences in sample size – for each trait across all adults captured at each site during each sampling period using the *cv* function in the ‘base’ R package. This resulted in six estimates for each trait from sites on our southern transect and four from sites on our northern transect. With these coefficients, we used linear mixed-effect models in the R package ‘lme4’ (Bates et al., 2015) to test whether the amount of phenotypic variation varied with elevation, where the coefficient of variation was the dependent variable, elevation was the predictor variable, and trait, year, and season were included as random effects. Because different types of traits might experience different selective regimes, we separately ran these linear models for each of three trait groups: metabolic traits (e.g., RMR, M_sum_, and metabolic scope); morphological traits (e.g., head length, head width, tarsus-metatarsus length, body mass, and body condition); and hematological traits (e.g., hemoglobin concentration and hematocrit).

Finally, to determine how patterns of survival and selection differed across elevations, we used a Bayesian state-space formulation of the Cormack-Jolly-Seber (CJS) survival model (Kéry & Schaub, 2012; Royle, 2008). These flexible models can accommodate the fact that we captured individuals on different dates and over different periods of time at each site. We constructed daily encounter histories for all adults, beginning with an individual’s first capture. However, a substantial number of recaptured individuals lost their PIT tag between seasons (*n*=33). Because we were able to identify individuals that had been previously caught *via* the presence of an ear clip, and because we had genetic samples for all individuals, we used the relatedness matrix generated in *vcftools* to identify all 33 of the individuals that had lost their tags and were recaptured at a later date. We then merged the encounter histories of these individuals from before and after their tag loss.

We built separate survival models for each of the 10 phenotypic traits to determine the strength of selection acting on each trait. Because many of the traits that we measured are likely seasonally flexible (Cheviron et al., 2013; Hayes, 1989), we standardized all variables separately for each site and sampling period (Gelman, 2008). Thus, we measured the strength of selection not on absolute trait values, but on site- and season-specific ones.

A trait’s influence on survival could be equivalent across all of our demographic sites, change systematically with elevation, or vary idiosyncratically across sites. We therefore modeled survival (□) as a logit linear function of either **(1)** transect, elevation, trait, and a trait*elevation interaction or **(2)** transect, site, trait, and a trait*site interaction. In both cases, transect was parameterized as a random effect on the intercept to account for the fact that survey sites were nested within separate elevational transects. Then, irrespective of whether we used elevation or site to model □, we used site and a measure of sampling effort – the number of trapping nights (3-7) at a site in a given season – to model detectability (α). We fit models using Markov chain Monte Carlo (MCMC) sampling implemented in JAGS using the R packages ‘rjags’ (Plummer, 2003, 2019) and ‘jagsUI’ (Kellner, 2024). We used uninformative priors for all parameters and sampled posterior distributions using three Markov chains of 25,000 iterations with a burn-in period of 1000 iterations and a thinning rate of 10. We assessed chain convergence by inspecting trace plots and calculating the Gelman-Rubin statistic (all Rhat values < 1.1). Finally, we report point estimates as posterior means in logit-form with 95% credible intervals and consider any trait for which we found that its 95% credible interval did not cross 0 to be under selection at either a particular site or across elevations. See the Supplemental Materials for more details.

## RESULTS

### Genetic Variation

Using 180 individuals from our six demographic sites, we found that pairwise F_st_ measures among sites ranged from 0.02-0.05. The degree of divergence consistently differed among sites along and between the two transects: pairwise F_st_ measures among sites along our southern transect were higher (x□ =0.04±0.001, *n*=5) than those between sites across our two transects (x□ =0.03±0.01, *n*=15) or those among sites on our northern transect (x□ =0.02±0.002, *n*=5; Supplemental Figure 1), but there was no relationship between F_st_ and the elevational difference between sites (β=1.37×10^-4^, SE=9.48×10^-5^, CI=-4.92×10^-5^, 3.22×10^-4^). Likewise, the site frequency spectra differed among sites (Supplemental Figure 2), with the D statistic from pairwise Kolmogorov-Smirnov tests between sites ranging from 0.03-0.10 (x =0.07±0.02, *n*=15). However, as the elevational difference between the two sites increased, the D statistic decreased (β=-2.91×10^-4^, SE=1.05×10^-4^, CI=-4.96×10^-4^, -8.54×10^-5^). Finally, measures of individual heterozygosity ranged between 0.25-0.34 (x□ =0.29±0.02, *n*=180) and trended lower at higher elevations (β=-1.20×10^-5^, SE=7.77×10^-6^, CI=-2.72×10^-5^, 3.24×10^-6^; Fig. 2), with the population at our highest elevation site having ∼1% less heterozygosity than the population at our lowest elevation site.

**Figure 2.**
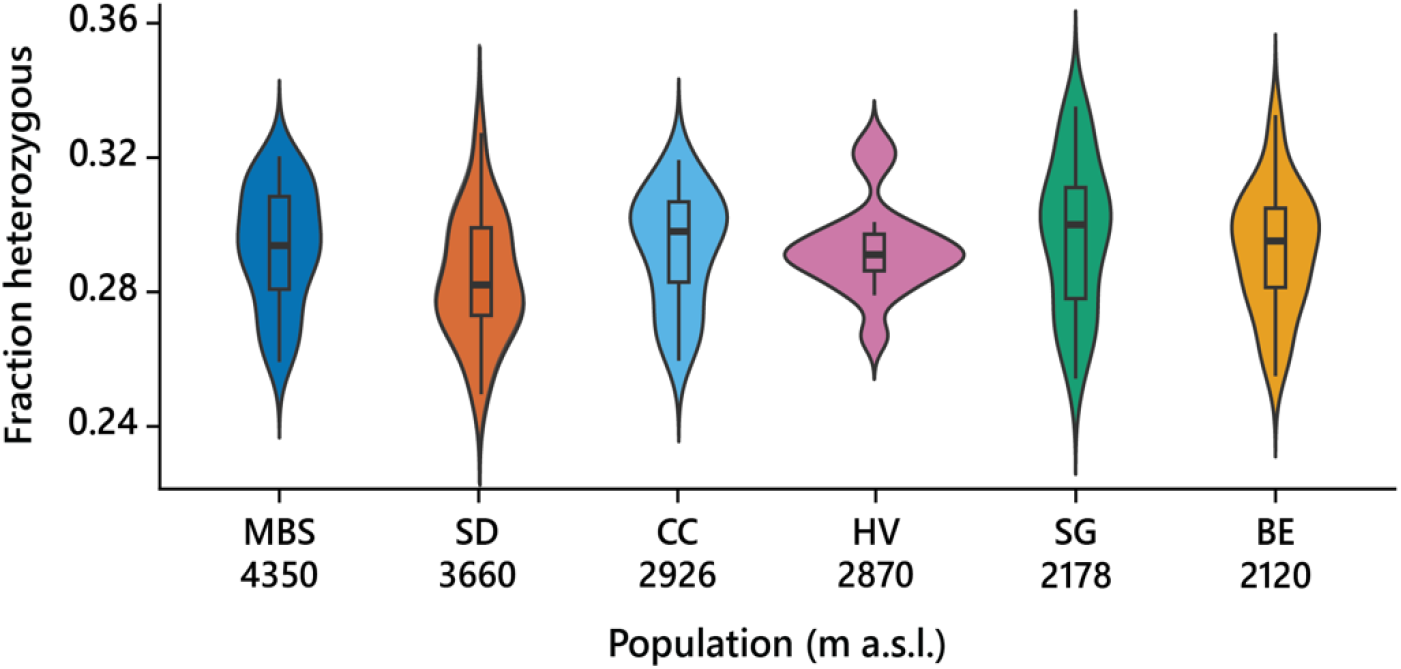
The fraction of sites across the genome at which individual deer mice (*Peromyscus maniculatus*) exhibited heterozygosity at six demographic plots along two elevational transects from 2,120-4,200m in the Rocky Mountains of Colorado, U.S.A.

### Phenotypic Variation

Across all phenotypic traits, variation declined as elevation increased, meaning that high elevation populations had ∼23% less trait variation than low elevation populations (β=-0.003, SE=0.001, CI=-0.005, -0.001; *n*=30 season × sites; Fig. 3). This pattern was clearest among metabolic traits (β=-0.003, SE=0.001, CI=-0.01, -0.001) and hematological traits (β=-0.002, SE=0.001, CI=-0.003, -0.001), but was less clear for morphological traits (β=-0.003, SE=0.002, CI=-0.006, 0.0003).

**Figure 3.**
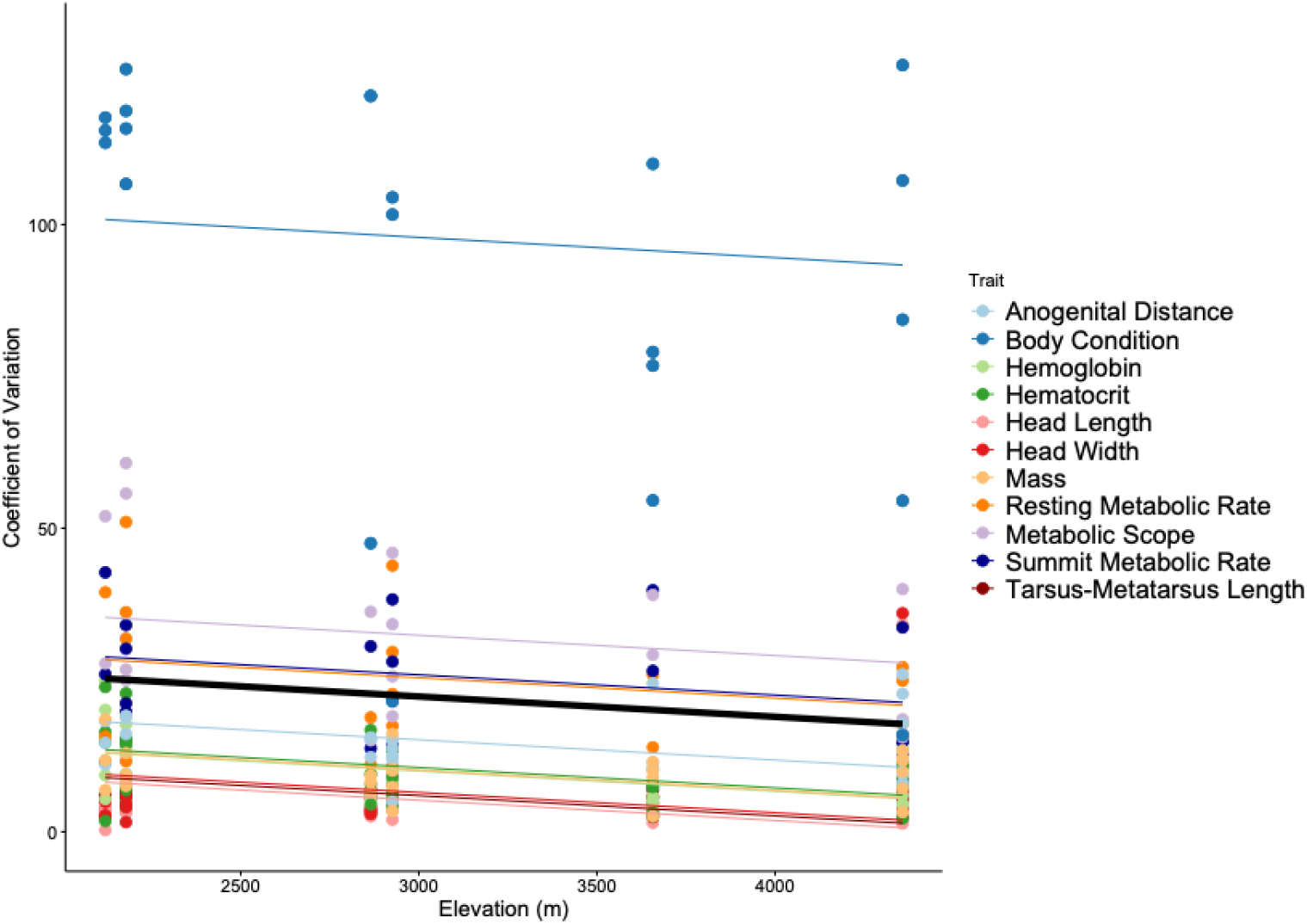
Relationship between elevation and phenotypic trait variation in deer mice (*Peromyscus maniculatus*) along two elevational transects ranging from 2120-4200 m in the Rocky Mountains of Colorado, U.S.A. Dots represent a site and year specific coefficient of variation for each trait; colors represent separate traits; the thick represents the mean regression line across all traits, while the thin, colored lines represent the trait-specific regression lines.

**Figure 4.**
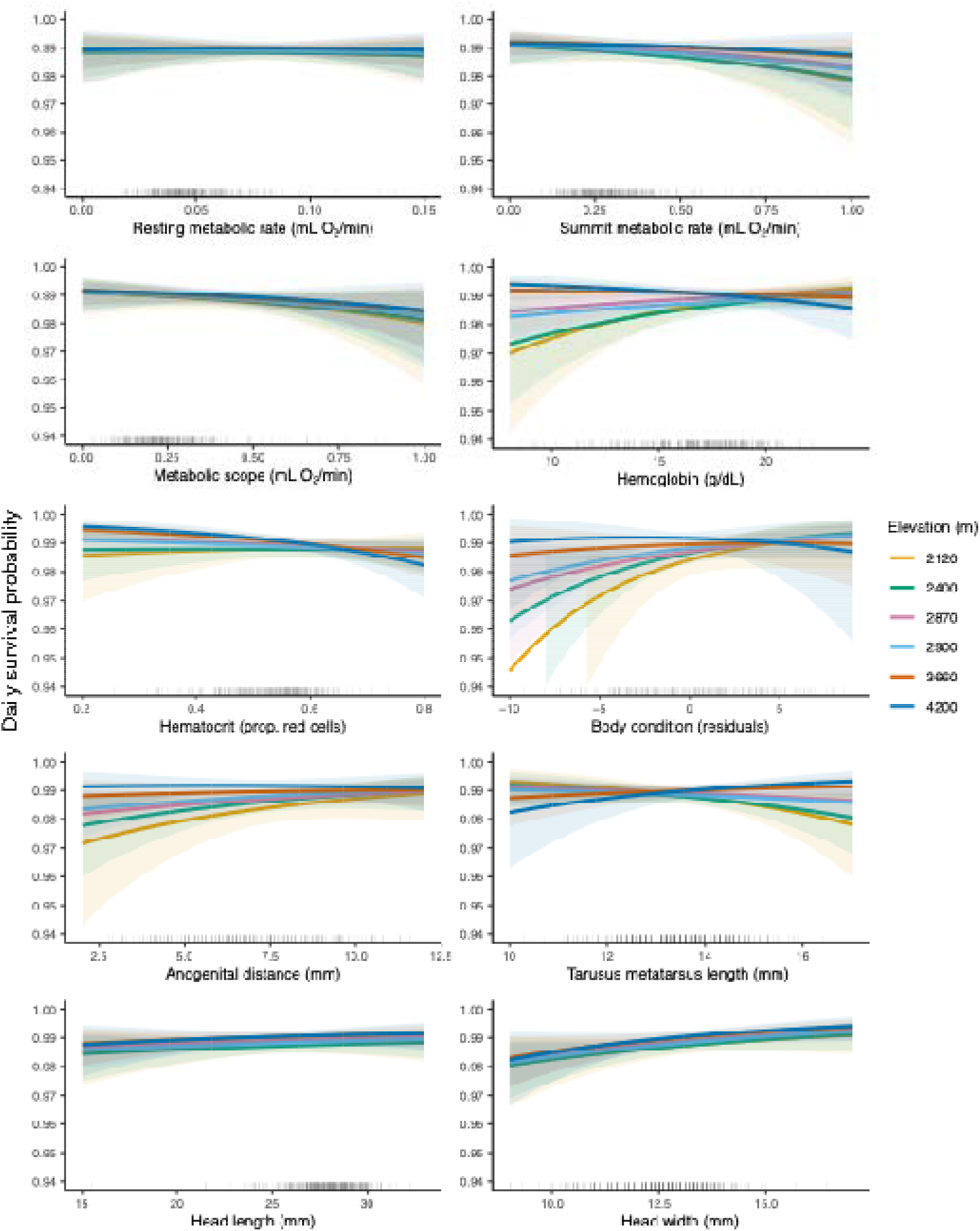
Relationship between phenotypic trait values and daily survival rates for deer mice (*Peromyscus maniculatus*) at six demographic sites along two elevational transects from 2,120-4,200m in the Rocky Mountains of Colorado, U.S.A. Colors represent the elevation of the separate demographic sites.

### The Influence of Traits and Elevation on Survival

The estimated strength of the relationship between survival and elevation varied between 0.11-0.45 (x =0.27±0.11; Table 1) across our 10 trait models. This variation in effect size was the result of differences in the sample size of individuals with trait measures for each respective trait. However, the models with the largest sample sizes (*n*=467 individuals; Table 1) had estimates with far less variance, ranging from 0.23-0.26 (x□ =0.25±0.01; Table 1). As such, mice at our highest elevation site were ∼7% more likely to survive for one month than mice from our lowest elevation site. Only one trait – head width – exhibited a consistent (positive) influence on survival across all elevations (β=0.30, SD=0.13, CI=0.06, 0.55). Likewise, only one trait – hemoglobin – exhibited a trend with elevation, whereby the strength of selection became stronger and more negative as elevation increased (elevation × trait: β=-0.50, SD=0.24, CI=-0.05, -0.96; Fig. 3). More than half of all traits had an influence on survival at one or more of our sites – body condition, metabolic scope, hemoglobin, hematocrit, head length, and head width. Finally, sites at 2870 and 2900m had the most traits under selection, with three and two traits that influenced survival, respectively (Supplementary Tables 2-11). Models with the largest sample sizes showed no difference in survival between transects (α_tran1_ = 4.46, SD= 0.07, CI = 4.32 – 4.73; α_tran2_ = 4.54, SD = 0.09, CI = 4.36 – 4.73), a result supported across all models.

**Table 1.** Results of Bayesian survival models investigating the influence of phenotypic trait values and elevation on the daily survival rates of deer mice (*Peromyscus maniculatus*) across two elevational transects from 2,120-4,200m in the Rocky Mountains of Colorado, U.S.A. Each trait was run in a model separately from the other traits. All estimates are on the logit scale and the 95% credible intervals around each model estimate are included in parentheses.

| **Trait** | **Main Trait Effect** | **Effect of Elevation** | **Trait × Elevation Interaction** | **Sample Size (Individuals)** |
|---|---|---|---|---|
| **Resting Metabolic Rate** | -0.03 (-0.29, 0.23) | 0.15 (-0.11, 0.41) | 0.04 (-0.50, 0.56) | 396 |
| **Summit Metabolic Rate** | -0.20 (-0.47, 0.08) | 0.27 (0.02, 0.,52) | 0.12 (-0.43, 0.66) | 388 |
| **Metabolic Scope** | -0.21 (-0.48, 0.08) | 0.11 (-0.15, 0.39) | 0.04 (-0.52, 0.60) | 357 |
| **Hematocrit** | -0.20 (-0.45, 0.05) | 0.20 (-0.06, 0.45) | -0.36 (-0.82, 0.11) | 402 |
| **Hemoglobin** | 0.11 (-0.14, 0.36) | 0.37 (0.12, 0.63) | -0.50 (-0.96, -0.05) | 380 |
| **Body Condition** | 0.16 (-0.10, 0.41) | 0.45 (0.13, 0.77) | -0.30 (-0.89, 0.28) | 260 |
| **Anogenital Distance** | 0.18 (-0.13, 0.51) | 0.44 (0.13, 0.76) | -0.29 (-0.94, 0.36) | 260 |
| **Head Length** | 0.11 (-0.12, 0.34) | 0.25 (0.01, 0.48) | 0.02 (-0.45, 0.49) | 467 |
| **Head Width** | 0.30 (0.06, 0.55) | 0.23 (0.002, 0.47) | 0.08 (-0.40, 0.56) | 467 |
| **Tarsus-Metatarsus** | -0.05 (-0.30, 0.18) | 0.25 (0.01, 0.49) | 0.44 (-0.05, 0.93) | 467 |

## DISCUSSION

Past studies have hypothesized that environmental gradients spanning from temperate to extreme environments should yield predictable evolutionary patterns, resulting in reductions in both phenotypic and genetic variation in more extreme environments (Conklin et al., 2017; Saether & Bakke, 2000). To test these predictions, we measured genetic variation, phenotypic trait variation, and survival in deer mice along two elevational transects ranging from 2,120-4,200m. In line with our predictions, we found that genetic and phenotypic variation both declined with elevation, especially in phenotypic traits related to blood oxygen carrying capacity and whole-organism aerobic performance. We also found that daily survival rates increased with elevation, as did the strength of selection on some hematological traits. Finally, we found that fewer traits influenced survival at high and low elevations than at middle elevations. Our results suggest that life in extreme environments can result in predictable patterns of evolution, which could help provide a basis for frameworks seeking to understand species’ vulnerabilities to environmental change.

### Extreme Environments and Trait Variation

Reductions in variation – whether genetic or phenotypic – are classic signatures of purifying selection. We thus predicted that, if higher elevation populations have experienced a history of strong, directional selection, we would observe reduced genetic and phenotypic variation in those populations relative to ones at lower elevations. Indeed, we found that, across all phenotypic traits, our highest elevation population had ∼23% less phenotypic variation than our lowest elevation population. We observed a similar trend – albeit not statistically significant – toward a reduction in heterozygosity in 818 SNPs spread across the genome in higher elevation populations. We likewise found a similar trend in heterozygosity (β=-2.98×10^-6^, SE=1.83×10^-6^, CI=-6.56×10^-6^, 5.99×10^-7^) in a *posthoc* analysis using data from an exome capture array (Schweizer et al. 2021) that was filtered similarly and included a subset of the individuals from this study (*n*=21 individuals), but many more sites across the genome (*n*=364,523).

Such patterns in trait variation could, however, have emerged from other, historical demographic processes. For instance, if the deer mouse populations colonizing high elevations were small and had little initial genetic variation, contemporary populations could still exhibit reduced levels of genetic diversity. Such founder effects would likely also result in variation in the site frequency spectra, with bottlenecked populations exhibiting right-skewed spectra – i.e., with few rare variants occurring across the genome (Freund et al., 2023; Lucena Perez et al., 2021). Instead, while we did observe differences in site frequency spectra among populations, the spectra of high and low elevation populations were largely similar, while the spectra of those populations differed more significantly from the spectra of populations at middle elevations. At the same time, our pairwise F_st_ measures indicated a degree of evolutionary independence among populations, even at similar elevations. Our results therefore point toward independent colonizations of high elevation environments by reasonably large founder populations followed by a history of shared selection pressures across elevations (see also, Schweizer et al., 2021).

### The Demographic Importance of Survival

Fitness in consistent and predictable environments should largely be determined by few, survival-related traits (Saether & Bakke, 2000). In turn, we predicted that survival rates would be higher at high elevations and that, at these elevations, there would only be selection on a small number of traits. We found that deer mice at higher elevations had 7% higher monthly survival rates than those at low elevations. We also found that few traits influenced survival at high elevations: both our elevation- and site-based models indicated that only hemoglobin concentration influenced survival at high elevations. Finally, and unexpectedly, we found that the largest number of traits were under selection at our two mid-elevation sites – not our lowest elevation sites – where six of the 10 traits we measured influenced survival.

It is telling that the trait we found to be under increasingly strong selection at higher elevations was hemoglobin concentration. Hemoglobin is key to an organism’s blood-oxygen carrying capacity and, as such, an important component of aerobic performance. The influence of the trait on aerobic performance at high elevations is, however, less pronounced than that of other traits that we were unable to measure (e.g., pulmonary O_2_ extraction and arterial O_2_ saturation; Tate et al., 2020). As such, hemoglobin concentration may be under contemporary selection because selection has already operated on other, more immediately consequential traits, and winnowed the variation in those traits to the point where there is population uniformity (see also, Velotta et al., 2025). Indeed, in this study, we found that high elevation mice exhibited relatively less variation in our whole-organism measure of aerobic performance – M_sum_ – than in hemoglobin concentration, while we have previously found genomic signatures of strong purifying selection on loci associated with M_sum_ at our highest elevation site (Corder, 2024). Our high elevation populations may therefore be approaching equilibrium with their environments.

But, why would our middle elevation sites have more traits under contemporary selection than our low elevation ones? It is possible that our elevational transects were characterized by more than one environmental gradient. Along our elevational transect, our lowest elevation sites are substantially more arid than our middle or high elevation ones (NOAA 2025). Our middle elevation sites could, therefore, counterintuitively exist at a point along multiple environmental gradients where selection pressures frequently shift from year-to-year, while our low and high elevation sites experience more predictably harsh conditions and a more limited range of selection pressures. If this were the case, however, we would not have expected phenotypic and genetic variation to decline with elevation, as both did in our study. Nonetheless, traits related to water conservation and heat dissipation (e.g., kidney morphology; Blumstein & MacManes, 2023) might be more likely to be under selection at our low elevation site than traits related to hypoxia and aerobic performance. Future studies should explore this possibility.

It is instead more likely that the fitness landscape is more varied and idiosyncratic at our middle and low elevation sites. This is to be expected if the factors imposing selection on these populations frequently fluctuate across time. Many small mammal populations experience regular population cycles over periods ranging from 7-10 years (Andreassen et al., 2021), meaning that studies must observe the entire range of potential population sizes and environmental conditions experienced in order to characterize the true drivers of fitness (Siepielski et al., 2009). Even during the length of this study, for example, mice at our low elevation sites experienced substantial variation in the prevalence of parasitic botflies (*Cuterebra* sp.), which are a known cause of mortality (Wilde et al., 2019). More time may be necessary to accurately characterize the fitness landscape at these elevations and in temperate environments more generally.

### Towards a Framework for Understanding Vulnerability to Environmental Change

Our results indicate that occupying environments close to the end of at least one environmental gradient can result in predictable evolutionary patterns for populations and thus, more generally, that environmental predictability is key to understanding species’ evolutionary histories. If this is indeed broadly true, our results can help contextualize the recent, rapid declines of other extremophile populations (e.g., Baker et al., 2004; Studds et al., 2017): those populations may also have had limited trait variation (but see Pegan et al., 2025), leaving them unable to respond to novel (unpredictable) environmental conditions (Conklin et al., 2017). In this light, efforts to identify where along environmental gradients tipping points exist between demographic predictability and seeming idiosyncrasy are critically important. So too are efforts to better characterize the full suite of relevant environmental gradients. Many gradients are already well studied (e.g., salinity; Whitehead et al., 2011), but there may remain hidden environmental heterogeneity waiting to be characterized but already predictably influencing populations’ evolutionary trajectories (Gulson-Castillo et al., 2023). Ultimately, it is by fully cataloging the types and amounts of environmental variation and predictability that populations face that we will be able to accurately forecast how they will fare in our increasingly unpredictable future.

## ACKNOWLEDGEMENTS

We are immensely grateful to the many field assistants who contributed to this project. We are also appreciative of the efforts of the staff of Rocky Mountain National Park and the Roosevelt and Arapaho National Forests for help with permitting. Finally, thank you to the member of the Senner Lab at the University of Massachusetts Amherst for helpful comments on an earlier version of this manuscript.

## AUTHOR CONTRIBUTIONS

NRS and ZAC conceived of the study; NRS, RM, KTS, RMS, JPV, LRW, CJW, and ZAC carried out the field work; NRS and KTS performed the lab work; NRS, BRA, TCN, RMS, SS, LRW, and MS contributed to the analyses; NRS wrote the manuscript, and all other co-authors offered feedback and edits.

## Supplementary Material

### METHODS

#### Respirometry Trials

We pumped ambient air through a darkened cabinet housing a Peltier device that regulated the temperature and kept the internal air at 27°C for RMR trials and -5°C for M_sum_ trials. We used mass flow controllers (Sable Systems Inc., Las Vegas, NV, USA) that maintained a flow rate of 500 ml/min for the in current ambient air during the RMR trials, while we pumped a heliox mixture (21% O2, 79% He ± 0.5%) equilibrated to the local atmospheric pressure at a flow rate of 750 ml/min in the M_sum_ trials. We scrubbed this air of H_2_O using drierite before it entered either the airtight baseline or animal respirometry chambers (polypropylene, 180 ml) and scrubbed it of CO_2_ with ascarite after it passed through those chambers. We then re dried it with drierite before pumping it through an oxygen analyzer (FoxBox, Sable Systems Inc.). For the M_sum_ trials, after a 5 min baseline, we monitored the excurrent O_2_ of a single mouse. We halted the trial when this individual’s oxygen consumption declined or it reached 30 minutes, at which point we collected a second, 5 min baseline. We used a rectal thermocouple to measure the body temperature of each individual before and after the trial to confirm it was hypothermic (e.g., an internal temperature ≤90% of the individual’s initial temperature) and verify it had reached M_sum_.

#### RadSeq Filtering Checks

To evaluate whether this filtering scheme was overly conservative, we compared diversity metrics from this dataset against a more relaxed alternative, retaining the same individuals and allele-frequency/depth filters but a more permissive site-completeness threshold. Observed heterozygosity and allele-frequency distributions were similar across the range of thresholds tested, but individuals admitted under looser individual-missingness cutoffs introduced a heterozygote deficit – inferred from a shift in the inbreeding coefficient, *F*, which switched from 0 to >0 – consistent with coverage- and missing-data-driven genotyping artifacts that can bias heterozygosity and related diversity estimates in RADseq data (O’Leary et al., 2018; Gautier et al., 2013). We additionally screened for cryptic relatedness using KING-robust kinship estimation in PLINK (v2.0; Manichaikul et al. 2010) and for linkage disequilibrium among our final SNP set; however, we did not remove related individuals or prune for linkage given the low portion of SNPs affected and the biological plausibility of localized kin structure among sampled mice.

#### Survival Models

CJS models include two components: a state process describing whether individual *i* is alive at time *t* that is unobserved or partially observed, and an observation process, which is conditional on the state process. We let the latent variable *z* denote the true state of an individual, where *z_i,t_* = 1 if it is alive and *z_i,t_*= 0 if it is dead. Individuals are alive at the occasion of their first capture, and states on subsequent occasions are modeled as Bernoulli trials, where, conditional on being alive at occasion *t* - 1, individual *i* survives to occasion *t* with probability □*_i,t_* _-1_. We modeled □*_i,t_* _-1_ as a logit linear function of transect, elevation, trait, and trait * elevation interaction:

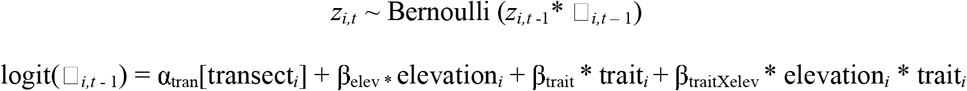

where α and β parameters are estimated regression coefficients. Transect was parameterized as a random effect on the intercept to account for the fact that survey sites were nested within elevational transects. The observation process was also modeled as realizations of Bernoulli trials, where individual *i* alive at occasion *t* is recaptured with probability *p_i,t_* _-1_. We modeled *p_i,t-1_* with separate intercepts for each site, and included an additive effect of survey effort to account for variation in the number of trapping nights (3-7) per site during spring and fall sampling periods over the course of the study:

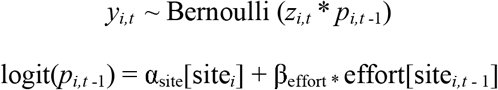

Including the latent variable *z* in the observation Bernoulli trials ensured that dead individuals could not be resighted. We fit models using Markov chain Monte Carlo (MCMC) sampling implemented in JAGS using the R packages *rjags* (Plummer 2003, 2019) and *jagsUI* (Kellner 2024). We used uninformative priors for all parameters and sampled posterior distributions using three Markov chains of 25,000 iterations with a burn-in period of 1000 iterations and a thinning rate of 10. We assessed chain convergence by inspecting trace plots and by calculating the Gelman-Rubin statistic (all Rhat values < 1.1). We report point estimates as posterior means in logit-form and 95% credible intervals.

### RESULTS

**Supplementary Table 1.** Step-by-step summary of the number of sites and individuals removed during our SNP filtering process.

| Step | Filter applied | Ind. before | Ind. after | Ind. removed | %Ind. removed | Sites before | Sites after | Sites removed | %Sites removed |
|---|---|---|---|---|---|---|---|---|---|
| Starting dataset | Removed 3 flagged (repeat) individuals | 489 | 486 | 3 | 0.6 | 2,183,359 | 2,183,359 | 0 | 0.0 |
| 1. Individual missingness | Removed individuals with >80% missing genotypes | 486 | 339 | 147 | 30.2 | 2,183,359 | 2,183,359 | 0 | 0.0 |
| 2. Allele frequency and depth | MAF ≥ 0.10; minimum mean site depth ≥ 5× | 339 | 339 | 0 | 0.0 | 2,183,359 | 9,225 | 2,174,134 | 99.6 |
| 3. Site missingness | Max-missing 50% (genotyped in ≥50% of individuals) | 339 | 339 | 0 | 0.0 | 9,225 | 9,044 | 181 | 2.0 |
| 4. Individual missingness | Removed individuals with >30% missing genotypes | 339 | 196 | 143 | 42.2 | 9,044 | 9,044 | 0 | 0.0 |
| 5. Site missingness | Max-missing 70% | 196 | 196 | 0 | 0.0 | 9,044 | 8,020 | 1,024 | 11.3 |
| 6. Individual missingness | Removed individuals with >25% missing genotypes | 196 | 180 | 16 | 8.2 | 8,020 | 8,020 | 0 | 0.0 |
| **7. Final Site missingness** | **Max-missing 95%** | **180** | **180** | **0** | **0.0** | **8,020** | **818** | **7,202** | **89.8** |

**Supplementary Table 2.** Results of a Bayesian survival model including a site x trait interaction for resting metabolic rate.

| **Parameter** | **Mean** | **St. Dev.** | **2.50%** | **97.50%** | **F** | **Rhat** | **# Eff** |
|---|---|---|---|---|---|---|---|
| **Survival** |  |  |  |  |  |  |  |
| Southern Transect | 4.51 | 0.07 | 4.37 | 4.65 | 1.00 | 1.00 | 7200 |
| Northern Transect | 4.50 | 0.10 | 4.31 | 4.69 | 1.00 | 1.00 | 5104 |
| Big Elk * RMR | -0.20 | 0.33 | -0.85 | 0.43 | 0.73 | 1.00 | 3416 |
| Spring Gulch * RMR | 0.05 | 0.32 | -0.58 | 0.69 | 0.55 | 1.00 | 7200 |
| Hidden Valley * RMR | -0.20 | 0.42 | -1.05 | 0.63 | 0.69 | 1.00 | 7200 |
| Chicago Creek * RMR | 0.14 | 0.26 | -0.36 | 0.65 | 0.71 | 1.00 | 5239 |
| Sundance Mountain * RMR | -0.13 | 0.35 | -0.82 | 0.56 | 0.65 | 1.00 | 3025 |
| Mount Blue Sky * RMR | -0.02 | 0.29 | -0.60 | 0.54 | 0.53 | 1.00 | 7200 |
| Big Elk | 1.48 | 0.29 | 0.94 | 2.06 | 1.00 | 1.00 | 2745 |
| Spring Gulch | -0.04 | 0.18 | -0.39 | 0.33 | 0.59 | 1.00 | 1804 |
| Hidden Valley | 1.17 | 0.19 | 0.81 | 1.57 | 1.00 | 1.00 | 7200 |
| Chicago Creek | 1.26 | 0.16 | 0.95 | 1.59 | 1.00 | 1.00 | 4784 |
| Sundance Mountain | 1.37 | 0.21 | 0.97 | 1.79 | 1.00 | 1.00 | 2672 |
| Mount Blue Sky | 0.24 | 0.13 | -0.01 | 0.50 | 0.97 | 1.00 | 3907 |
| **Detectability** |  |  |  |  |  |  |  |
| Effort | -0.26 | 0.20 | -0.66 | 0.14 | 0.90 | 1.00 | 3046 |
| **Deviance** | 1800.91 | 26.42 | 1750.65 | 1853.09 | 1.00 | 1.00 | 893 |

**Supplementary Table 3.** Results of a Bayesian survival model including a site x trait interaction for summit metabolic rate.

| **Parameter** | **Mean** | **St. Dev.** | **2.50%** | **97.50%** | **F** | **Rhat** | **# Eff** |
|---|---|---|---|---|---|---|---|
| **Survival** |  |  |  |  |  |  |  |
| Southern Transect | 4.48 | 0.08 | 4.33 | 4.63 | 1.00 | 1.00 | 2265 |
| Northern Transect | 4.52 | 0.10 | 4.33 | 4.72 | 1.00 | 1.00 | 7200 |
| Big Elk * M<sub>sum</sub> | 0.00 | 0.43 | -0.85 | 0.84 | 0.50 | 1.00 | 7200 |
| Spring Gulch * M<sub>sum</sub> | -0.28 | 0.29 | -0.85 | 0.30 | 0.83 | 1.00 | 7200 |
| Hidden Valley * M<sub>sum</sub> | 0.19 | 0.49 | -0.76 | 1.16 | 0.65 | 1.00 | 7200 |
| Chicago Creek * M<sub>sum</sub> | -0.61 | 0.31 | -1.22 | 0.01 | 0.97 | 1.00 | 7200 |
| Sundance Mountain * M<sub>sum</sub> | 0.27 | 0.44 | -0.60 | 1.12 | 0.74 | 1.00 | 7200 |
| Mount Blue Sky * M<sub>sum</sub> | -0.17 | 0.27 | -0.72 | 0.33 | 0.75 | 1.00 | 3389 |
| Big Elk | 1.47 | 0.30 | 0.91 | 2.07 | 1.00 | 1.00 | 2125 |
| Spring Gulch | 0.27 | 0.15 | -0.03 | 0.57 | 0.96 | 1.00 | 3221 |
| Hidden Valley | 1.11 | 0.19 | 0.74 | 1.49 | 1.00 | 1.00 | 7200 |
| Chicago Creek | 1.25 | 0.17 | 0.92 | 1.60 | 1.00 | 1.00 | 3610 |
| Sundance Mountain | 1.39 | 0.21 | 0.99 | 1.82 | 1.00 | 1.00 | 7200 |
| Mount Blue Sky | 0.14 | 0.12 | -0.10 | 0.39 | 0.88 | 1.00 | 7200 |
| **Detectability** |  |  |  |  |  |  |  |
| Effort | -0.02 | 0.18 | -0.38 | 0.34 | 0.54 | 1.00 | 6519 |
| **Deviance** | 1857.54 | 26.16 | 1807.24 | 1910.36 | 1.00 | 1.00 | 7200 |

**Supplementary Table 4.** Results of a Bayesian survival model including a site x trait interaction for metabolic scope.

| **Parameter** | **Mean** | **St. Dev.** | **2.50%** | **97.50%** | **F** | **Rhat** | **# Eff** |
|---|---|---|---|---|---|---|---|
| **Survival** |  |  |  |  |  |  |  |
| Southern Transect | 4.48 | 0.08 | 4.32 | 4.64 | 1.00 | 1.00 | 5914 |
| Northern Transect | 4.47 | 0.10 | 4.27 | 4.67 | 1.00 | 1.00 | 7200 |
| Big Elk * Metabolic Scope | 0.01 | 0.44 | -0.85 | 0.88 | 0.51 | 1.00 | 7200 |
| Spring Gulch * Metabolic Scope | -0.27 | 0.32 | -0.91 | 0.36 | 0.80 | 1.00 | 7200 |
| Hidden Valley * Metabolic Scope | 0.24 | 0.49 | -0.73 | 1.22 | 0.69 | 1.00 | 7200 |
| Chicago Creek * Metabolic Scope | -0.60 | 0.31 | -1.24 | -0.01 | 0.98 | 1.00 | 7200 |
| Sundance Mountain * Metabolic Scope | 0.32 | 0.47 | -0.59 | 1.26 | 0.75 | 1.00 | 7200 |
| Mount Blue Sky * Metabolic Scope | -0.24 | 0.28 | -0.79 | 0.29 | 0.81 | 1.00 | 7200 |
| Big Elk | 1.46 | 0.29 | 0.92 | 2.06 | 1.00 | 1.00 | 7200 |
| Spring Gulch | -0.03 | 0.18 | -0.39 | 0.33 | 0.57 | 1.00 | 7200 |
| Hidden Valley | 1.17 | 0.19 | 0.81 | 1.56 | 1.00 | 1.00 | 7200 |
| Chicago Creek | 1.23 | 0.17 | 0.91 | 1.57 | 1.00 | 1.00 | 5175 |
| Sundance Mountain | 1.38 | 0.23 | 0.95 | 1.85 | 1.00 | 1.00 | 7200 |
| Mount Blue Sky | 0.18 | 0.13 | -0.08 | 0.45 | 0.91 | 1.00 | 7200 |
| **Detectability** |  |  |  |  |  |  |  |
| Effort | -0.13 | 0.21 | -0.55 | 0.29 | 0.73 | 1.00 | 7200 |
| **Deviance** | 1646.72 | 24.83 | 1598.90 | 1695.22 | 1.00 | 1.00 | 7200 |

**Supplementary Table 5.** Results of a Bayesian survival model including a site x trait interaction for hemoglobin concentration.

| **Parameter** | **Mean** | **St. Dev.** | **2.50%** | **97.50%** | **F** | **Rhat** | **# Eff** |
|---|---|---|---|---|---|---|---|
| **Survival** |  |  |  |  |  |  |  |
| Southern Transect | 4.45 | 0.08 | 4.30 | 4.61 | 1.00 | 1.00 | 3420 |
| Northern Transect | 4.51 | 0.10 | 4.33 | 4.70 | 1.00 | 1.00 | 1086 |
| Big Elk * Hemoglobin | 0.31 | 0.35 | -0.36 | 1.01 | 0.81 | 1.00 | 4735 |
| Spring Gulch * Hemoglobin | 0.18 | 0.28 | -0.37 | 0.75 | 0.73 | 1.00 | 7200 |
| Hidden Valley * Hemoglobin | 0.23 | 0.45 | -0.65 | 1.13 | 0.69 | 1.00 | 7200 |
| Chicago Creek * Hemoglobin | 0.31 | 0.31 | -0.27 | 0.95 | 0.85 | 1.00 | 6114 |
| Sundance Mountain * Hemoglobin | 0.44 | 0.35 | -0.23 | 1.13 | 0.90 | 1.00 | 7200 |
| Mount Blue Sky * Hemoglobin | -0.46 | 0.24 | -0.93 | -0.01 | 0.98 | 1.00 | 7200 |
| Big Elk | 1.21 | 0.25 | 0.74 | 1.72 | 1.00 | 1.00 | 7200 |
| Spring Gulch | 0.37 | 0.17 | 0.04 | 0.70 | 0.99 | 1.00 | 1269 |
| Hidden Valley | 1.02 | 0.18 | 0.67 | 1.38 | 1.00 | 1.00 | 7200 |
| Chicago Creek | 1.07 | 0.19 | 0.71 | 1.44 | 1.00 | 1.00 | 7200 |
| Sundance Mountain | 1.26 | 0.19 | 0.90 | 1.64 | 1.00 | 1.00 | 7200 |
| Mount Blue Sky | 0.25 | 0.12 | 0.01 | 0.49 | 0.98 | 1.00 | 3668 |
| **Detectability** |  |  |  |  |  |  |  |
| Effort | 0.04 | 0.17 | -0.30 | 0.38 | 0.59 | 1.00 | 838 |
| **Deviance** | 1855.85 | 27.39 | 1801.49 | 1910.67 | 1.00 | 1.00 | 7200 |

**Supplemental Table 6.** Results of a Bayesian survival model including a site x trait interaction for hematocrit.

| **Parameter** | **Mean** | **St. Dev.** | **2.50%** | **97.50%** | **F** | **Rhat** | **# Eff** |
|---|---|---|---|---|---|---|---|
| **Survival** |  |  |  |  |  |  |  |
| Southern Transect | 4.50 | 0.08 | 4.35 | 4.65 | 1.00 | 1.00 | 7200 |
| Northern Transect | 4.52 | 0.09 | 4.34 | 4.71 | 1.00 | 1.00 | 7200 |
| Big Elk * Hematocrit | 0.30 | 0.34 | -0.37 | 0.96 | 0.81 | 1.00 | 7200 |
| Spring Gulch * Hematocrit | -0.20 | 0.34 | -0.87 | 0.47 | 0.72 | 1.00 | 7200 |
| Hidden Valley * Hematocrit | 0.54 | 0.42 | -0.26 | 1.39 | 0.90 | 1.00 | 2825 |
| Chicago Creek * Hematocrit | -0.70 | 0.31 | -1.33 | -0.11 | 0.99 | 1.00 | 4691 |
| Sundance Mountain * Hematocrit | -0.10 | 0.36 | -0.81 | 0.62 | 0.61 | 1.00 | 7200 |
| Mount Blue Sky * Hematocrit | -0.46 | 0.21 | -0.88 | -0.05 | 0.99 | 1.00 | 7200 |
| Big Elk | 1.18 | 0.25 | 0.71 | 1.70 | 1.00 | 1.00 | 2096 |
| Spring Gulch | 0.23 | 0.19 | -0.14 | 0.59 | 0.89 | 1.00 | 6538 |
| Hidden Valley | 1.04 | 0.18 | 0.70 | 1.41 | 1.00 | 1.00 | 2373 |
| Chicago Creek | 1.35 | 0.16 | 1.05 | 1.68 | 1.00 | 1.00 | 7200 |
| Sundance Mountain | 1.30 | 0.19 | 0.94 | 1.68 | 1.00 | 1.00 | 7200 |
| Mount Blue Sky | 0.24 | 0.12 | 0.01 | 0.49 | 0.98 | 1.00 | 7200 |
| **Detectability** |  |  |  |  |  |  |  |
| Effort | -0.14 | 0.18 | -0.48 | 0.20 | 0.79 | 1.00 | 6982 |
| **Deviance** | 1891.92 | 28.34 | 1836.70 | 1947.48 | 1.00 | 1.00 | 5207 |

**Supplemental Table 7.** Results of a Bayesian survival model including a site x trait interaction for body condition (males only).

| **Parameter** | **Mean** | **St. Dev.** | **2.50%** | **97.50%** | **F** | **Rhat** | **# Eff** |
|---|---|---|---|---|---|---|---|
| **Survival** |  |  |  |  |  |  |  |
| Southern Transect | 4.46 | 0.07 | 4.33 | 4.60 | 1.00 | 1.00 | 1207 |
| Northern Transect | 4.47 | 0.09 | 4.29 | 4.66 | 1.00 | 1.00 | 3955 |
| Big Elk * Body Condition | 0.38 | 0.30 | -0.17 | 0.99 | 0.91 | 1.00 | 5425 |
| Spring Gulch * Body Condition | -0.02 | 0.26 | -0.53 | 0.48 | 0.52 | 1.00 | 3198 |
| Hidden Valley * Body Condition | 1.15 | 0.43 | 0.35 | 2.02 | 1.00 | 1.00 | 2720 |
| Chicago Creek * Body Condition | 0.63 | 0.28 | 0.09 | 1.19 | 0.99 | 1.00 | 4422 |
| Sundance Mountain * Body Condition | -0.47 | 0.37 | -1.17 | 0.25 | 0.90 | 1.00 | 7200 |
| Mount Blue Sky * Body Condition | -0.20 | 0.27 | -0.74 | 0.34 | 0.77 | 1.00 | 7025 |
| Big Elk | 1.14 | 0.24 | 0.68 | 1.63 | 1.00 | 1.00 | 7200 |
| Spring Gulch | 0.25 | 0.15 | -0.04 | 0.54 | 0.95 | 1.00 | 6461 |
| Hidden Valley | 1.09 | 0.18 | 0.74 | 1.45 | 1.00 | 1.00 | 7200 |
| Chicago Creek | 1.32 | 0.16 | 1.01 | 1.64 | 1.00 | 1.00 | 7200 |
| Sundance Mountain | 1.22 | 0.18 | 0.86 | 1.58 | 1.00 | 1.00 | 2312 |
| Mount Blue Sky | 0.19 | 0.12 | -0.04 | 0.42 | 0.95 | 1.00 | 7200 |
| **Detectability** |  |  |  |  |  |  |  |
| Effort | -0.02 | 0.17 | -0.35 | 0.30 | 0.55 | 1.00 | 7200 |
| **Deviance** | 2139.24 | 28.75 | 2084.26 | 2196.14 | 1.00 | 1.00 | 7200 |

**Supplemental Table 8.** Results of a Bayesian survival model including a site x trait interaction for anogenital distance (males only).

| **Parameter** | **Mean** | **St. Dev.** | **2.50%** | **97.50%** | **F** | **Rhat** | **# Eff** |
|---|---|---|---|---|---|---|---|
| **Survival** |  |  |  |  |  |  |  |
| Southern Transect | 4.48 | 0.07 | 4.35 | 4.61 | 1.00 | 1.00 | 7200 |
| Northern Transect | 4.52 | 0.09 | 4.34 | 4.71 | 1.00 | 1.00 | 7200 |
| Big Elk * Anogenital Distance | -0.43 | 0.33 | -1.06 | 0.20 | 0.91 | 1.00 | 3668 |
| Spring Gulch * Anogenital Distance | -0.06 | 0.26 | -0.57 | 0.45 | 0.60 | 1.00 | 7200 |
| Hidden Valley * Anogenital Distance | 0.19 | 0.34 | -0.49 | 0.85 | 0.71 | 1.00 | 7200 |
| Chicago Creek * Anogenital Distance | 0.05 | 0.23 | -0.39 | 0.49 | 0.58 | 1.00 | 7200 |
| Sundance Mountain * Anogenital Distance | -0.21 | 0.29 | -0.79 | 0.33 | 0.77 | 1.00 | 7200 |
| Mount Blue Sky * Anogenital Distance | 0.21 | 0.23 | -0.24 | 0.65 | 0.82 | 1.00 | 7200 |
| Big Elk | 1.14 | 0.25 | 0.69 | 1.64 | 1.00 | 1.00 | 7200 |
| Spring Gulch | 0.24 | 0.15 | -0.04 | 0.53 | 0.95 | 1.00 | 3420 |
| Hidden Valley | 1.07 | 0.18 | 0.72 | 1.43 | 1.00 | 1.00 | 7200 |
| Chicago Creek | 1.32 | 0.16 | 1.02 | 1.65 | 1.00 | 1.00 | 7200 |
| Sundance Mountain | 1.21 | 0.19 | 0.87 | 1.58 | 1.00 | 1.00 | 4501 |
| Mount Blue Sky | 0.19 | 0.12 | -0.04 | 0.43 | 0.95 | 1.00 | 7200 |
| **Detectability** |  |  |  |  |  |  |  |
| Effort | -0.03 | 0.17 | -0.35 | 0.29 | 0.57 | 1.00 | 7200 |
| **Deviance** | 2150.62 | 29.34 | 2094.54 | 2208.75 | 1.00 | 1.00 | 3602 |

**Supplemental Table 9.** Results of a Bayesian survival model including a site x trait interaction for tarsal-metatarsal length.

| **Parameter** | **Mean** | **St. Dev.** | **2.50%** | **97.50%** | **F** | **Rhat** | **# Eff** |
|---|---|---|---|---|---|---|---|
| **Survival** |  |  |  |  |  |  |  |
| Southern Transect | 4.47 | 0.07 | 4.34 | 4.60 | 1.00 | 1.00 | 4115 |
| Northern Transect | 4.50 | 0.09 | 4.32 | 4.69 | 1.00 | 1.00 | 5463 |
| Big Elk * Tarsus-Metatarsus | -0.66 | 0.41 | -1.51 | 0.08 | 0.96 | 1.00 | 7200 |
| Spring Gulch * Tarsus-Metatarsus | 0.13 | 0.29 | -0.45 | 0.71 | 0.66 | 1.00 | 4088 |
| Hidden Valley * Tarsus-Metatarsus | -0.50 | 0.44 | -1.35 | 0.36 | 0.88 | 1.00 | 7200 |
| Chicago Creek * Tarsus-Metatarsus | -0.10 | 0.25 | -0.61 | 0.39 | 0.65 | 1.00 | 7200 |
| Sundance Mountain * Tarsus-Metatarsus | -0.07 | 0.29 | -0.66 | 0.50 | 0.59 | 1.00 | 7200 |
| Mount Blue Sky * Tarsus-Metatarsus | 0.39 | 0.25 | -0.11 | 0.88 | 0.94 | 1.00 | 4835 |
| Big Elk | 1.14 | 0.24 | 0.68 | 1.64 | 1.00 | 1.00 | 7200 |
| Spring Gulch | 0.24 | 0.15 | -0.05 | 0.53 | 0.95 | 1.00 | 5337 |
| Hidden Valley | 1.06 | 0.18 | 0.70 | 1.43 | 1.00 | 1.00 | 7200 |
| Chicago Creek | 1.32 | 0.16 | 1.01 | 1.64 | 1.00 | 1.00 | 5348 |
| Sundance Mountain | 1.21 | 0.19 | 0.86 | 1.58 | 1.00 | 1.00 | 7200 |
| Mount Blue Sky | 0.19 | 0.12 | -0.04 | 0.43 | 0.95 | 1.00 | 2742 |
| **Detectability** |  |  |  |  |  |  |  |
| Effort | -0.03 | 0.17 | -0.35 | 0.30 | 0.56 | 1.00 | 1781 |
| **Deviance** | 2147.37 | 28.73 | 2091.12 | 2203.50 | 1.00 | 1.00 | 1848 |

**Supplemental Table 10.** Results of a Bayesian survival model including a site x trait interaction for head length.

| **Parameter** | **Mean** | **St. Dev.** | **2.50%** | **97.50%** | **F** | **Rhat** | **# Eff** |
|---|---|---|---|---|---|---|---|
| **Survival** |  |  |  |  |  |  |  |
| Southern Transect | 4.48 | 0.07 | 4.34 | 4.61 | 1.00 | 1.00 | 7200 |
| Northern Transect | 4.48 | 0.09 | 4.30 | 4.67 | 1.00 | 1.00 | 7200 |
| Big Elk * Head Length | -0.12 | 0.32 | -0.75 | 0.51 | 0.64 | 1.00 | 7200 |
| Spring Gulch * Head Length | -0.05 | 0.26 | -0.55 | 0.46 | 0.58 | 1.00 | 7200 |
| Hidden Valley * Head Length | 1.16 | 0.47 | 0.29 | 2.09 | 1.00 | 1.00 | 7200 |
| Chicago Creek * Head Length | 0.29 | 0.30 | -0.30 | 0.88 | 0.84 | 1.00 | 7200 |
| Sundance Mountain * Head Length | -0.17 | 0.29 | -0.75 | 0.40 | 0.72 | 1.00 | 7200 |
| Mount Blue Sky * Head Length | 0.11 | 0.27 | -0.42 | 0.63 | 0.66 | 1.00 | 7200 |
| Big Elk | 1.14 | 0.25 | 0.68 | 1.65 | 1.00 | 1.00 | 7200 |
| Spring Gulch | 0.25 | 0.15 | -0.05 | 0.54 | 0.95 | 1.00 | 5092 |
| Hidden Valley | 1.08 | 0.19 | 0.72 | 1.45 | 1.00 | 1.00 | 3385 |
| Chicago Creek | 1.32 | 0.16 | 1.01 | 1.64 | 1.00 | 1.00 | 7200 |
| Sundance Mountain | 1.22 | 0.18 | 0.87 | 1.58 | 1.00 | 1.00 | 2356 |
| Mount Blue Sky | 0.19 | 0.12 | -0.04 | 0.42 | 0.94 | 1.00 | 7200 |
| **Detectability** |  |  |  |  |  |  |  |
| Effort | -0.02 | 0.17 | -0.35 | 0.30 | 0.55 | 1.00 | 7200 |
| **Deviance** | 2146.47 | 29.64 | 2089.64 | 2205.43 | 1.00 | 1.00 | 7135 |

**Supplemental Table 11.** Results of a Bayesian survival model including a site x trait interaction for head width.

| **Parameter** | **Mean** | **St. Dev.** | **2.50%** | **97.50%** | **F** | **Rhat** | **# Eff** |
|---|---|---|---|---|---|---|---|
| **Survival** |  |  |  |  |  |  |  |
| Southern Transect | 4.47 | 0.07 | 4.34 | 4.60 | 1.00 | 1.00 | 5419 |
| Northern Transect | 4.47 | 0.10 | 4.28 | 4.66 | 1.00 | 1.00 | 7200 |
| Big Elk * Head Width | 0.54 | 0.35 | -0.12 | 1.23 | 0.95 | 1.00 | 3577 |
| Spring Gulch * Head Width | 0.05 | 0.27 | -0.48 | 0.55 | 0.57 | 1.00 | 2779 |
| Hidden Valley * Head Width | 1.03 | 0.47 | 0.15 | 1.99 | 0.99 | 1.00 | 5681 |
| Chicago Creek * Head Width | -0.05 | 0.28 | -0.60 | 0.50 | 0.58 | 1.00 | 7200 |
| Sundance Mountain * Head Width | 0.56 | 0.37 | -0.16 | 1.28 | 0.94 | 1.00 | 7200 |
| Mount Blue Sky * Head Width | 0.39 | 0.26 | -0.12 | 0.89 | 0.94 | 1.00 | 7200 |
| Big Elk | 1.16 | 0.25 | 0.69 | 1.66 | 1.00 | 1.00 | 1832 |
| Spring Gulch | 0.24 | 0.15 | -0.04 | 0.53 | 0.95 | 1.00 | 7200 |
| Hidden Valley | 1.07 | 0.18 | 0.73 | 1.43 | 1.00 | 1.00 | 7200 |
| Chicago Creek | 1.32 | 0.16 | 1.01 | 1.64 | 1.00 | 1.00 | 7200 |
| Sundance Mountain | 1.22 | 0.19 | 0.87 | 1.60 | 1.00 | 1.00 | 7200 |
| Mount Blue Sky | 0.19 | 0.12 | -0.04 | 0.41 | 0.95 | 1.00 | 2687 |
| **Detectability** |  |  |  |  |  |  |  |
| Effort | -0.03 | 0.17 | -0.35 | 0.30 | 0.58 | 1.00 | 7200 |
| **Deviance** | 2141.48 | 29.14 | 2085.52 | 2199.93 | 1.00 | 1.00 | 6243 |

**Supplemental Figure 1.**
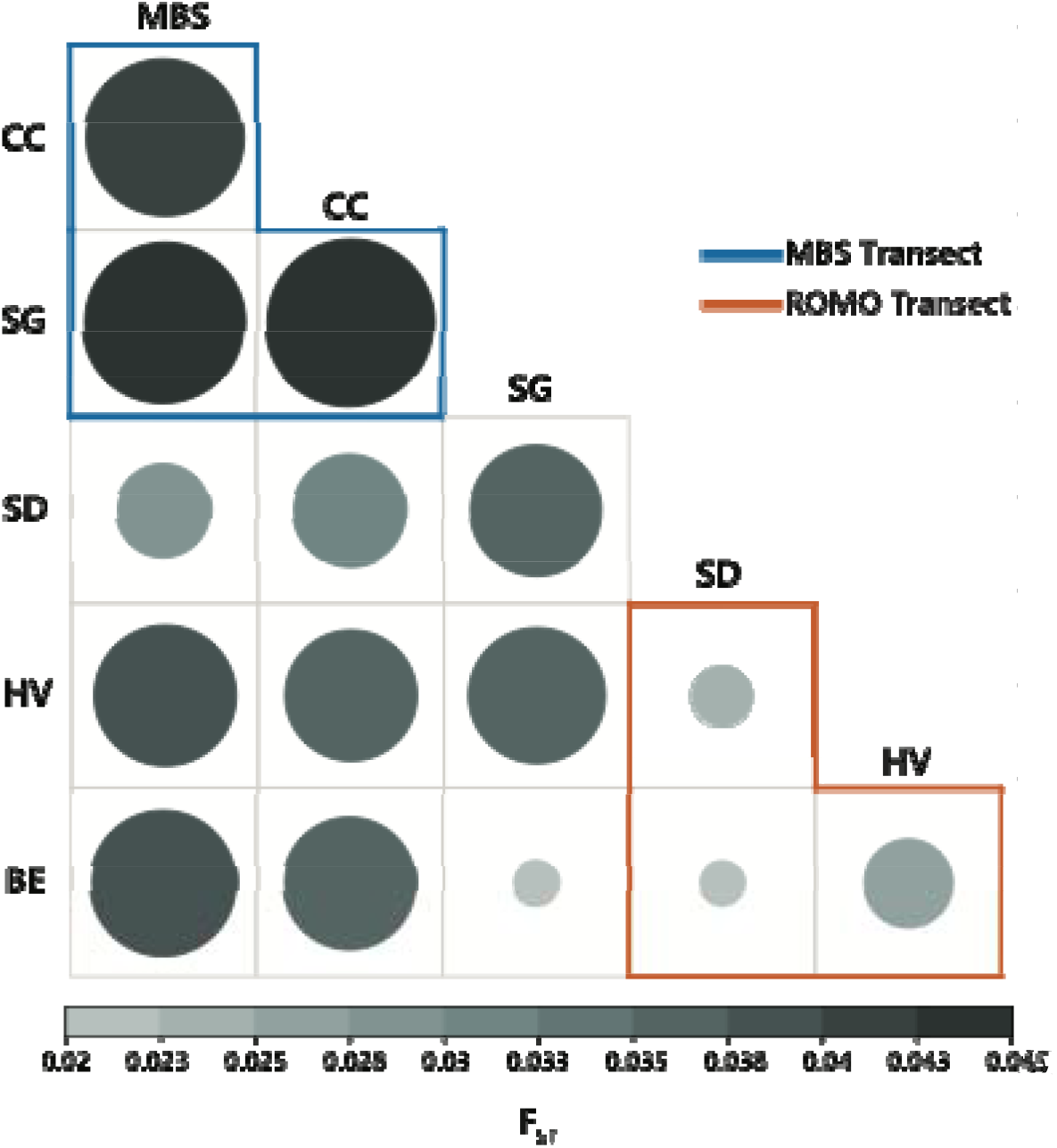
Pairwise comparisons of F_st_ among the six demographic plots. Sites from the two separate elevational transects are outlined in blue (the southern transect) or orange (northern transect). The size and color of the circle reflect the degree of divergence between the pair of sites.

**Supplemental Figure 2.**
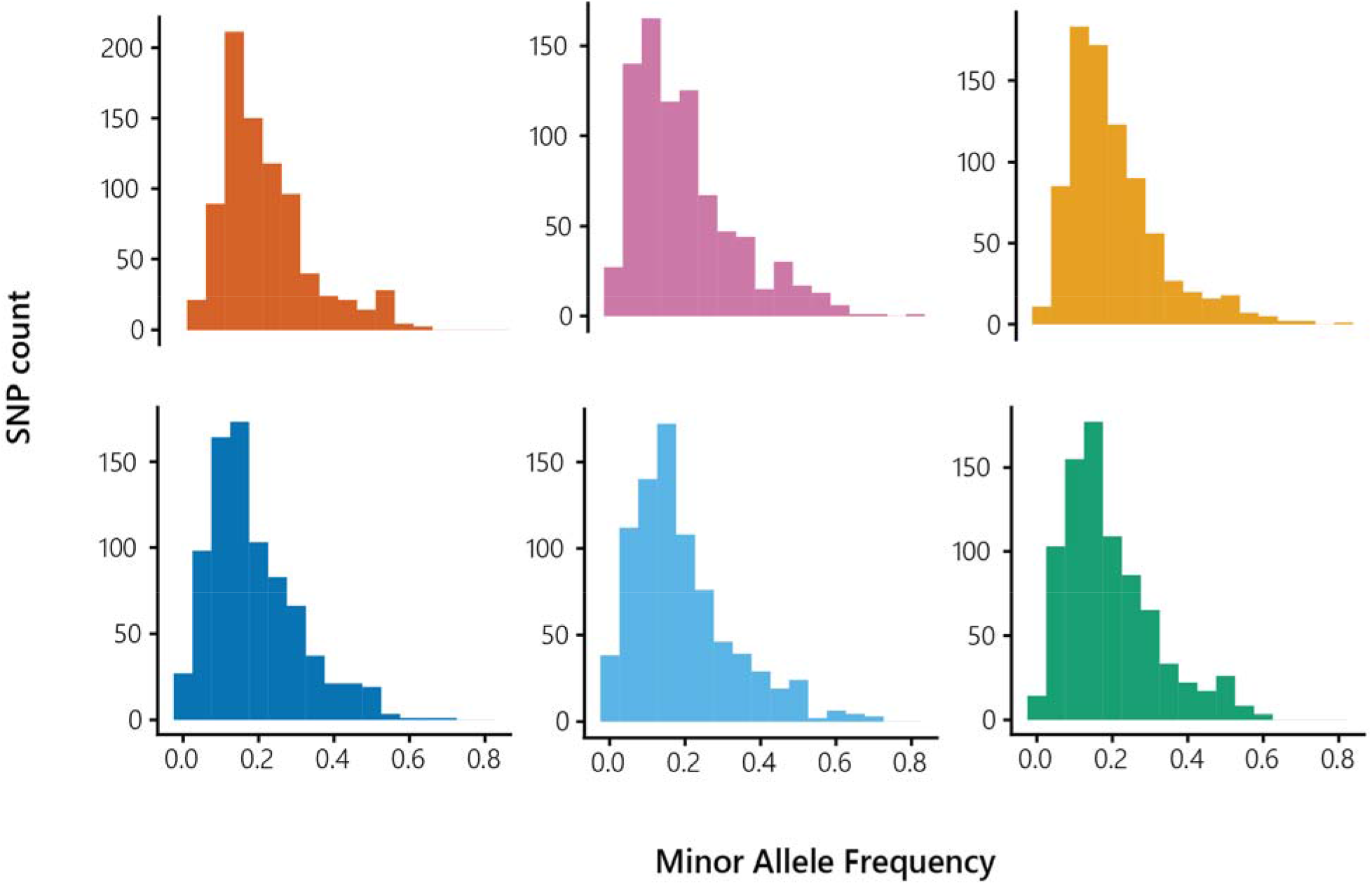
The site frequency spectrum from of the six demographic plot. Each demographic plot is represented by a separate panel. Within each panel, the height of each bar represents the count of single nucleotide polymorphisms (SNPs) with a particular minor allele frequency.

## REFERENCES

Ali, O. A., O’Rourke, S. M., Amish, S. J., Meek, M. H., Luikart, G., Jeffres, C., & Miller, M. R. (2016). RAD Capture (Rapture): Flexible and Efficient Sequence-Based Genotyping. Genetics, 202(2), 389–400. 10.1534/genetics.115.183665

Andreassen, H. P., Sundell, J., Ecke, F., Halle, S., Haapakoski, M., Henttonen, H., Huitu, O., Jacob, J., Johnsen, K., Koskela, E., Luque-Larena, J. J., Lecomte, N., Leirs, H., Mariën, J., Neby, M., Rätti, O., Sievert, T., Singleton, G. R., Van Cann, J., … Ylönen, H. (2021). Population cycles and outbreaks of small rodents: Ten essential questions we still need to solve. Oecologia, 195(3), 601–622. 10.1007/s00442-020-04810-w

Andrew, J. R., Saltzman, W., Chappell, M. A., & Garland, T. (2016). Consequences of Fatherhood in the Biparental California Mouse (Peromyscus californicus): Locomotor Performance, Metabolic Rate, and Organ Masses. Physiological & Biochemical Zoology, 89(2), 130–140. 10.1086/685435

Arnold, T., B., & Emerson, J., W. (2011). Nonparametric Goodness-of-Fit Tests for Discrete Null Distributions. The R Journal, 3(2), 34. 10.32614/RJ-2011-016

Baker, A. J., González, P. M., Piersma, T., Niles, L. J., de Lima Serrano do Nascimento, I., Atkinson, P. W., Clark, N. A., Minton, C. D. T., Peck, M. K., & Aarts, G. (2004). Rapid population decline in red knots: Fitness consequences of decreased refuelling rates and late arrival in Delaware Bay. Proceedings of the Royal Society of London. Series B: Biological Sciences, 271(1541), 875–882. 10.1098/rspb.2003.2663

Barrett, R. D. H., Laurent, S., Mallarino, R., Pfeifer, S. P., Xu, C. C. Y., Foll, M., Wakamatsu, K., Duke-Cohan, J. S., Jensen, J. D., & Hoekstra, H. E. (2018). The fitness consequences of genetic variation in wild populations of mice [Preprint]. Evolutionary Biology. 10.1101/383240

Bates, D., Mächler, M., Bolker, B., & Walker, S. (2015). Fitting Linear Mixed-Effects Models Using lme4. Journal of Statistical Software, 67(1). 10.18637/jss.v067.i01

Bertness, M. D., & Callaway, R. (1994). Positive interactions in communities. Trends in Ecology & Evolution, 9(5), 191–193. 10.1016/0169-5347(94)90088-4

Bitter, M. C., Wong, J. M., Dam, H. G., Donelan, S. C., Kenkel, C. D., Komoroske, L. M., Nickols, K. J., Rivest, E. B., Salinas, S., Burgess, S. C., & Lotterhos, K. E. (2021). Fluctuating selection and global change: A synthesis and review on disentangling the roles of climate amplitude, predictability and novelty. Proceedings of the Royal Society B: Biological Sciences, 288(1957), 20210727. 10.1098/rspb.2021.0727

Blumstein, D. M., & MacManes, M. D. (2023). When the tap runs dry: The physiological effects of acute experimental dehydration in Peromyscus eremicus. Journal of Experimental Biology, 226(23), jeb246386. 10.1242/jeb.246386

Botero, C. A., Weissing, F. J., Wright, J., & Rubenstein, D. R. (2015). Evolutionary tipping points in the capacity to adapt to environmental change. Proceedings of the National Academy of Sciences, 112(1), 184–189. 10.1073/pnas.1408589111

Cahill, A. E., Aiello-Lammens, M. E., Fisher-Reid, M. C., Hua, X., Karanewsky, C. J., Yeong Ryu, H., Sbeglia, G. C., Spagnolo, F., Waldron, J. B., Warsi, O., & Wiens, J. J. (2013). How does climate change cause extinction? Proceedings of the Royal Society B: Biological Sciences, 280(1750), 20121890. 10.1098/rspb.2012.1890

Catchen, J., Hohenlohe, P. A., Bassham, S., Amores, A., & Cresko, W. A. (2013). Stacks: An analysis tool set for population genomics. Molecular Ecology, 22(11), 3124–3140. 10.1111/mec.12354

Catchen, J. M., Amores, A., Hohenlohe, P., Cresko, W., & Postlethwait, J. H. (2011). Stacks: Building and Genotyping Loci De Novo From Short-Read Sequences. G3 Genes|Genomes|Genetics, 1(3), 171–182. 10.1534/g3.111.000240

Chappell, M. A., Hammond, K. A., Cardullo, R. A., Russell, G. A., Rezende, E. L., & Miller, C. (2007). Deer Mouse Aerobic Performance across Altitudes: Effects of Developmental History and Temperature Acclimation. Physiological and Biochemical Zoology, 80(6), 652–662. 10.1086/521202

Cheviron, Z. A., Bachman, G. C., Connaty, A. D., McClelland, G. B., & Storz, J. F. (2012). Regulatory changes contribute to the adaptive enhancement of thermogenic capacity in high-altitude deer mice. Proceedings of the National Academy of Sciences, 109(22), 8635–8640. 10.1073/pnas.1120523109

Cheviron, Z. A., Bachman, G. C., & Storz, J. F. (2013). Contributions of phenotypic plasticity to differences in thermogenic performance between highland and lowland deer mice. Journal of Experimental Biology, 216(7), 1160–1166. 10.1242/jeb.075598

Cheviron, Z. A., Connaty, A. D., McClelland, G. B., & Storz, J. F. (2014). FUNCTIONAL GENOMICS OF ADAPTATION TO HYPOXIC COLD-STRESS IN HIGH-ALTITUDE DEER MICE: TRANSCRIPTOMIC PLASTICITY AND THERMOGENIC PERFORMANCE: SPECIAL SECTION. Evolution, 68(1), 48–62. 10.1111/evo.12257

Cheviron, Z. A., Natarajan, C., Projecto-Garcia, J., Eddy, D. K., Jones, J., Carling, M. D., Witt, C. C., Moriyama, H., Weber, R. E., Fago, A., & Storz, J. F. (2014). Integrating Evolutionary and Functional Tests of Adaptive Hypotheses: A Case Study of Altitudinal Differentiation in Hemoglobin Function in an Andean Sparrow, Zonotrichia capensis. Molecular Biology and Evolution, 31(11), 2948–2962. 10.1093/molbev/msu234

Conklin, J. R., Senner, N. R., Battley, P. F., & Piersma, T. (2017). Extreme migration and the individual quality spectrum. Journal of Avian Biology, 48(1), 19–36. 10.1111/jav.01316

Corder, K. R. (2024). GENOMIC AND PHENOTYPIC INSIGHTS INTO THE MECHANISMS OF ADAPTATION IN HIGH-ELEVATION DEER MICE.

Djawdan, M., & Garland, T. (1988). Maximal Running Speeds of Bipedal and Quadrupedal Rodents. Journal of Mammalogy, 69(4), 765–772. 10.2307/1381631

Freund, F., Kerdoncuff, E., Matuszewski, S., Lapierre, M., Hildebrandt, M., Jensen, J. D., Ferretti, L., Lambert, A., Sackton, T. B., & Achaz, G. (2023). Interpreting the pervasive observation of U-shaped Site Frequency Spectra. PLOS Genetics, 19(3), e1010677. 10.1371/journal.pgen.1010677

Gabriel, W., Luttbeg, B., Sih, A., & Tollrian, R. (2005). Environmental tolerance, heterogeneity, and the evolution of reversible plastic responses. The American Naturalist, 166(3), 339–353. 10.1086/432558

Gelman, A. (2008). Scaling regression inputs by dividing by two standard deviations. Statistics in Medicine, 27(15), 2865–2873. 10.1002/sim.3107

Gerstein, A. C., Lo, D. S., & Otto, S. P. (2012). Parallel Genetic Changes and Nonparallel Gene–Environment Interactions Characterize the Evolution of Drug Resistance in Yeast. Genetics, 192(1), 241–252. 10.1534/genetics.112.142620

Gerstein, A. C., Ono, J., Lo, D. S., Campbell, M. L., Kuzmin, A., & Otto, S. P. (2015). Too Much of a Good Thing: The Unique and Repeated Paths Toward Copper Adaptation. Genetics, 199(2), 555–571. 10.1534/genetics.114.171124

Godsall, B., Coulson, T., & Malo, A. F. (2014). From physiology to space use: Energy reserves and androgenization explain home-range size variation in a woodland rodent. Journal of Animal Ecology, 83(1), 126–135. 10.1111/1365-2656.12116

Greenway, R., Barts, N., Henpita, C., Brown, A. P., Arias-Rodriguez, L., Rodríguez Peña, C. M., Arndt, S., Lau, G. Y., Murphy, M. P., Wu, L., Lin, D., Shaw, J. H., Kelley, J. L., & Tobler, M. (2020). Convergent evolution of conserved mitochondrial pathways underlies repeated adaptation to extreme environments [Preprint]. Evolutionary Biology. 10.1101/2020.02.24.959916

Gulson-Castillo, E. R., Van Doren, B. M., Bui, M. X., Horton, K. G., Li, J., Moldwin, M. B., Shedden, K., Welling, D. T., & Winger, B. M. (2023). Space weather disrupts nocturnal bird migration. Proceedings of the National Academy of Sciences, 120(42), e2306317120. 10.1073/pnas.2306317120

Hayes, J. P. (1989). Altitudinal and seasonal effects on aerobic metabolism of deer mice. Journal of Comparative Physiology B, 159(4), 453–459. 10.1007/BF00692417

Hayes, J. P., & O’Connor, C. S. (1999). Natural Selection on Thermogenic Capacity of High-Altitude Deer Mice. Evolution, 53(4), 1280. 10.2307/2640830

Hock, R. J. (1964). Physiological responses of deer mice to various native altitudes. In The Physiological Effects of High Altitude (pp. 59–72). Pergamon.

Ickin, E., Conquet, E., Abrahms, B., Albon, S., Blumstein, D., Bond, M., Boersma, P., Clark-Wolf, T., Clutton-Brock, T., Compagnoni, A., Dostálek, T., Evers, S., Fichtel, C., Gamelon, M., García-Callejas, D., Griesser, M., Hansen, B., Jenouvrier, S., Jerstad, K., … Paniw, M. (2024). Comparative Life-Cycle Analyses Reveal Interacting Climatic and Biotic Drivers of Population Responses to Climate Change. Life Sciences. 10.32942/X24C92

Kellner, K. (2024). jagsUI: A wrapper around “rjags” to streamline “JAGS” analyses. R package version 1.6.2. Https://CRAN.R-project.org/package=jagsUI [Computer software].

Kéry, M., & Schaub, M. (2012). Estimation of survival from capture-recapture data using the Cormack-Jolly-Seber model. In Bayesian population analysis uisng WinBUGS (pp. 172–237). Academic Press.

Kolis, K. M., Berg, C. S., Nelson, T. C., & Fishman, L. (2022). Population genomic consequences of life-history and mating system adaptation to a geothermal soil mosaic in yellow monkeyflowers. Evolution, 76(4), 765–781. 10.1111/evo.14469

Lanfear, R., Kokko, H., & Eyre-Walker, A. (2014). Population size and the rate of evolution. Trends in Ecology & Evolution, 29(1), 33–41. 10.1016/j.tree.2013.09.009

Lau, D. S., Connaty, A. D., Mahalingam, S., Wall, N., Cheviron, Z. A., Storz, J. F., Scott, G. R., & McClelland, G. B. (2017). Acclimation to hypoxia increases carbohydrate use during exercise in high-altitude deer mice. American Journal of Physiology-Regulatory, Integrative and Comparative Physiology, 312(3), R400–R411. 10.1152/ajpregu.00365.2016

Leiby, N., & Marx, C. J. (2014). Metabolic erosion primarily through mutation accumulation, and not tradeoffs, drives limited evolution of substrate specificity in Escherichia coli. PLoS Biology, 12(2), e1001789. 10.1371/journal.pbio.1001789

Li, H., & Durbin, R. (2009). Fast and accurate short read alignment with Burrows–Wheeler transform. Bioinformatics, 25(14), 1754–1760. 10.1093/bioinformatics/btp324

Li, H., Handsaker, B., Wysoker, A., Fennell, T., Ruan, J., Homer, N., Marth, G., Abecasis, G., Durbin, R., & 1000 Genome Project Data Processing Subgroup. (2009). The Sequence Alignment/Map format and SAMtools. Bioinformatics, 25(16), 2078–2079. 10.1093/bioinformatics/btp352

Lighton, J. R. B. (2008). Measuring metabolic rates: A manual for scientists. Oxford University Press.

Losos, J. B., & Ricklefs, R. E. (2009). Adaptation and diversification on islands. Nature, 457(7231), Article 7231. 10.1038/nature07893

Lucena-Perez, M., Kleinman-Ruiz, D., Marmesat, E., Saveljev, A. P., Schmidt, K., & Godoy, J. A. (2021). Bottleneck-associated changes in the genomic landscape of genetic diversity in wild lynx populations. Evolutionary Applications, 14(11), 2664–2679. 10.1111/eva.13302

Lui, M. A., Mahalingam, S., Patel, P., Connaty, A. D., Ivy, C. M., Cheviron, Z. A., Storz, J. F., McClelland, G. B., & Scott, G. R. (2015). High-altitude ancestry and hypoxia acclimation have distinct effects on exercise capacity and muscle phenotype in deer mice. American Journal of Physiology-Regulatory, Integrative and Comparative Physiology, 308(9), R779–R791. 10.1152/ajpregu.00362.2014

Malekar, V. C., Morton, J. D., Hider, R. N., Cruickshank, R. H., Hodge, S., & Metcalf, V. J. (2018). Effect of elevated temperature on membrane lipid saturation in Antarctic notothenioid fish. PeerJ, 6, e4765. 10.7717/peerj.4765

Mas, A., Lagadeuc, Y., & Vandenkoornhuyse, P. (2020). Reflections on the Predictability of Evolution: Toward a Conceptual Framework. iScience, 23(11). 10.1016/j.isci.2020.101736

Pegan, T. M., Kimmitt, A. A., Benz, B. W., Weeks, B. C., Aubry, Y., Burg, T. M., Hudon, J., Jones, A. W., Kirchman, J. J., Ruegg, K. C., & Winger, B. M. (2025). Long-distance seasonal migration to the tropics promotes genetic diversity but not gene flow in boreal birds. Nature Ecology & Evolution, 9(6), 957–969. 10.1038/s41559-025-02699-3

Pfeifer, S. P., Laurent, S., Sousa, V. C., Linnen, C. R., Foll, M., Excoffier, L., Hoekstra, H. E., & Jensen, J. D. (2017). The evolutionary history of Nebraska deer mice: Local adaptation in the face of strong gene flow [Preprint]. Evolutionary Biology. 10.1101/152694

Plummer, M. (2003). JAGS: a program for analysis of Bayesian graphical models using Gibbs sampling. Proceedings of the 3rd International Workshop on Distributed Statistical Computing, 125, 1–10.

Plummer, M. (2019). Rjags: Bayesian graphical models using MCMC. R package version 4.14. Https://CRAN.R-project.org/package=rjags [Computer software].

Reiskind, M. O. B., Moody, M. L., Bolnick, D. I., Hanifin, C. T., & Farrior, C. E. (2021). Nothing in Evolution Makes Sense Except in the Light of Biology. BioScience, (biaa170). 10.1093/biosci/biaa170

Royle, J. A. (2008). Modeling individual effects in the Cormack-Jolly-Seber model: A state-space formulation. Biometrics, 64(2), 364–370.

Saether, B.-E., & Bakke, O. (2000). Avian Life History Variation and Contribution of Demographic Traits to the Population Growth Rate. Ecology, 81, 642–653.

Schweizer, R. M., Jones, M. R., Bradburd, G. S., Storz, J. F., Senner, N. R., Wolf, C., & Cheviron, Z. A. (2021). Broad concordance in the spatial distribution of adaptive and neutral genetic variation across an elevational gradient in deer mice. Molecular Biology and Evolution, msab161. 10.1093/molbev/msab161

Schweizer, R. M., Velotta, J. P., Ivy, C. M., Jones, M. R., Muir, S. M., Bradburd, G. S., Storz, J. F., Scott, G. R., & Cheviron, Z. A. (2019). Physiological and genomic evidence that selection on the transcription factor Epas1 has altered cardiovascular function in high-altitude deer mice. PLOS Genetics, 15(11), e1008420. 10.1371/journal.pgen.1008420

Scott, G. R., Elogio, T. S., Lui, M. A., Storz, J. F., & Cheviron, Z. A. (2015). Adaptive Modifications of Muscle Phenotype in High-Altitude Deer Mice Are Associated with Evolved Changes in Gene Regulation. Molecular Biology and Evolution, 32(8), 1962–1976. 10.1093/molbev/msv076

Siepielski, A. M., DiBattista, J. D., & Carlson, S. M. (2009). It’s about time: The temporal dynamics of phenotypic selection in the wild. Ecology Letters, 12(11), 1261–1276. 10.1111/j.1461-0248.2009.01381.x

Smith, C. C., & Fretwell, S. D. (1974). The optimal balance between size and number of offspring. The American Naturalist, 108(962), 499–506.

Somero, G. N. (2010). The physiology of climate change: How potentials for acclimatization and genetic adaptation will determine “winners” and “losers.” Journal of Experimental Biology, 213(6), 912–920. 10.1242/jeb.037473

Speakman, J. R., Ergon, T., Cavanagh, R., Reid, K., Scantlebury, D. M., & Lambin, X. (2003). Resting and daily energy expenditures of free-living field voles are positively correlated but reflect extrinsic rather than intrinsic effects. Proceedings of the National Academy of Sciences, 100(24), 14057–14062. 10.1073/pnas.2235671100

Stager, M., Senner, N. R., Tobalske, B. W., & Cheviron, Z. A. (2020). Body temperature maintenance acclimates in a winter-tenacious songbird. The Journal of Experimental Biology, 223(12), jeb221853. 10.1242/jeb.221853

Stager, M., Velotta, J. P., Cheviron, Z. A., & Senner, N. R. (2025). Assessing the evidence for treating developmental plasticity and phenotypic flexibility as different phenomena. Functional Ecology, 39(11), 2949–2961. 10.1111/1365-2435.14590

Stasinopoulos, M. (2024). Gamlss (Version 5.4-22) [R]. doi:10.1111/j.1467-9876.2005.00510.x

Stern, D. L., & Orgogozo, V. (2008). The Loci of Evolution: How Predictable Is Genetic Evolution? Evolution, 62(9), 2155–2177.

Stevens, R. D., & Tello, J. S. (2018). A latitudinal gradient in dimensionality of biodiversity. Ecography, 41(12), 2016–2026. 10.1111/ecog.03654

Storz, J. F., Cheviron, Z. A., McClelland, G. B., & Scott, G. R. (2019). Evolution of physiological performance capacities and environmental adaptation: Insights from high-elevation deer mice (Peromyscus maniculatus). Journal of Mammalogy, 100(3), 910–922. 10.1093/jmammal/gyy173

Storz, J. F., Natarajan, C., Cheviron, Z. A., Hoffmann, F. G., & Kelly, J. K. (2012). Altitudinal Variation at Duplicated β-Globin Genes in Deer Mice: Effects of Selection, Recombination, and Gene Conversion. Genetics, 190(1), 203–216. 10.1534/genetics.111.134494

Storz, J. F., Runck, A. M., Sabatino, S. J., Kelly, J. K., Ferrand, N., Moriyama, H., Weber, R. E., & Fago, A. (2009). Evolutionary and functional insights into the mechanism underlying high-altitude adaptation of deer mouse hemoglobin. Proceedings of the National Academy of Sciences, 106(34), 14450–14455. 10.1073/pnas.0905224106

Storz, J. F., & Scott, G. R. (2019). Life Ascending: Mechanism and Process in Physiological Adaptation to High-Altitude Hypoxia. Annual Review of Ecology, Evolution, and Systematics, 50(1), 503–526. 10.1146/annurev-ecolsys-110218-025014

Storz, J. F., Scott, G. R., & Cheviron, Z. A. (2010). Phenotypic plasticity and genetic adaptation to high-altitude hypoxia in vertebrates. Journal of Experimental Biology, 213(24), 4125–4136. 10.1242/jeb.048181

Studds, C. E., Kendall, B. E., Murray, N. J., Wilson, H. B., Rogers, D. I., Clemens, R. S., Gosbell, K., Hassell, C. J., Jessop, R., Melville, D. S., Milton, D. A., Minton, C. D. T., Possingham, H. P., Riegen, A. C., Straw, P., Woehler, E. J., & Fuller, R. A. (2017). Rapid population decline in migratory shorebirds relying on Yellow Sea tidal mudflats as stopover sites. Nature Communications, 8(1), 14895. 10.1038/ncomms14895

Szűcs, M., Melbourne, B. A., Tuff, T., Weiss-Lehman, C., & Hufbauer, R. A. (2017). Genetic and demographic founder effects have long-term fitness consequences for colonising populations. Ecology Letters, 20(4), 436–444. 10.1111/ele.12743

Taborsky, B., English, S., Fawcett, T. W., Kuijper, B., Leimar, O., McNamara, J. M., Ruuskanen, S., & Sandi, C. (2021). Towards an Evolutionary Theory of Stress Responses. Trends in Ecology & Evolution, 36(1), 39–48. 10.1016/j.tree.2020.09.003

Tate, K. B., Ivy, C. M., Velotta, J. P., Storz, J. F., McClelland, G. B., Cheviron, Z. A., & Scott, G. R. (2017). Circulatory mechanisms underlying adaptive increases in thermogenic capacity in high-altitude deer mice. The Journal of Experimental Biology, 220(20), 3616–3620. 10.1242/jeb.164491

Tate, K. B., Wearing, O. H., Ivy, C. M., Cheviron, Z. A., Storz, J. F., McClelland, G. B., & Scott, G. R. (2020). Coordinated changes across the O2 transport pathway underlie adaptive increases in thermogenic capacity in high-altitude deer mice. Proceedings of the Royal Society B: Biological Sciences, 287(1927), 20192750. 10.1098/rspb.2019.2750

Tobler, M., Palacios, M., Chapman, L. J., Mitrofanov, I., Bierbach, D., Plath, M., Arias-Rodriguez, L., León, F. J. G. de, & Mateos, M. (2011). Evolution in Extreme Environments: Replicated Phenotypic Differentiation in Livebearing Fish Inhabiting Sulfidic Springs. Evolution, 65(8), 2213–2228. 10.1111/j.1558-5646.2011.01298.x

Velotta, J. P., Ivy, C. M., Wolf, C. J., Scott, G. R., & Cheviron, Z. A. (2018). Maladaptive phenotypic plasticity in cardiac muscle growth is suppressed in high-altitude deer mice. Evolution, 72(12), 2712–2727. 10.1111/evo.13626

Velotta, J. P., Stager, M., Cheviron, Z. A., & Senner, N. R. (2025). Incomplete genetic compensation and countergradient variation of blood-oxygen transport in deer mice. Evolution, 79(11), 2552–2562. 10.1093/evolut/qpaf186

Whitehead, A., Galvez, F., Zhang, S., Williams, L. M., & Oleksiak, M. F. (2011). Functional Genomics of Physiological Plasticity and Local Adaptation in Killifish. Journal of Heredity, 102(5), 499–511. 10.1093/jhered/esq077

Wiens, J. J., & Graham, C. H. (2005). Niche Conservatism: Integrating Evolution, Ecology, and Conservation Biology. Annual Review of Ecology, Evolution, and Systematics, 36(1), 519–539. 10.1146/annurev.ecolsys.36.102803.095431

Wilde, L. R., Wolf, C. J., Porter, S. M., Stager, M., Cheviron, Z. A., & Senner, N. R. (2019). Botfly infections impair the aerobic performance and survival of montane populations of deer mice, Peromyscus maniculatus rufinus. Functional Ecology, 33(4), 608–618. 10.1111/1365-2435.13276

Williams, O. (1955). Distribution of Mice and Shrews in a Colorado Montane Forest. Journal of Mammalogy, 36(2), 221. 10.2307/1375880

Williams, S. H., Peiffer, E., & Ford, S. (2009). Gape and bite force in the rodents Onychomys leucogaster and Peromyscus maniculatus: Does jaw-muscle anatomy predict performance? Journal of Morphology, 270(11), 1338–1347. 10.1002/jmor.10761

Wolf, C. J. (2023). VARIATION IN BODY COMPOSITION IN DEER MICE: THE BIOGEOGRAPHY AND ECOPHYSIOLOGY OF ENERGY ALLOCATION.

Zhang, X.-S. (2012). FISHER’S GEOMETRICAL MODEL OF FITNESS LANDSCAPE AND VARIANCE IN FITNESS WITHIN A CHANGING ENVIRONMENT: GENETIC VARIATION IN FITNESS. Evolution, 66(8), 2350–2368. 10.1111/j.1558-5646.2012.01610.x

